# Beyond Invariable Sites: Using Evolutionary Stasis to Map Multi-Layered Constraints on the Evolution of Viral and Mammalian Genomes

**DOI:** 10.64898/2026.04.09.717527

**Authors:** Sergei L. Kosakovsky Pond, Hannah Verdonk, Steven Weaver, Gallean Brown, Danielle Callan, Anton Nekrutenko, Darren P. Martin

**Affiliations:** Department of Biology, Temple University, Philadelphia, PA, USA; Institute of Infectious Disease and Molecular Virology, Division of Medical Virology, University of Cape Town, Cape Town, South Africa; Department of Biochemistry and Molecular Biology, The Pennsylvania State University, University Park, PA, USA

**Keywords:** Comparative Genomics, Codon Models, Purifying Selection, Evolutionary Stasis, B-STILL

## Abstract

The quantification of genomic conservation has progressed from foundational statistical modeling of evolutionary rates to state-of-the-art phylogeny-aware deep learning architectures. Yet, a fundamental resolution gap remains whenever evolutionary rates closely approach the “zero-rate origin,” where standard selection inference tools will essentially ignore signals of extreme purifying section at invariant genome sites. We present B-STILL (Bayesian Significance Test of Invariant Low Likelihoods), a hierarchical Bayesian framework designed to resolve the selective landscape of protein-coding data by leveraging gene-level calibration and codon-site specific evolutionary opportunity. This framework is based on computationally efficient approximations using codon-substitution models which are scalable to alignments with thousands of sequences. By explicitly tuning the stasis radius around the near-zero evolutionary-rate regime, B-STILL distinguishes between stochastic invariance and functional constraint, identifying Evolutionary Stasis Anchors (ESAs) where the upper bound on permitted evolutionary change is statistically anomalous relative to the background of the gene. This hierarchical approach provides a signature of functional or structural constraint that is often difficult to detect using other tools. Validation against extensive pathogen and clinical databases confirms that ESAs are predictors of biological fitness and disease potential. Collectively, we identified thousands of significantly clustered ESAs that precisely footprint both known functional domains and currently uncharacterized structural motifs in mammalian and viral genomes. These findings establish B-STILL as a scalable statistical framework for high-resolution genomic annotation, transforming formerly ignored invariant genome and protein sites into informative markers of extreme purifying selection across both well-characterized and uncharacterized protein-coding genes from different domains of life.

## Introduction

Historically, invariable nucleotide sites have been viewed as uninformative for phylogenetic reconstruction, as positions lacking variation provide no signal for resolving tree topology. Consequently, these sites were often relegated to background covariates—most notably the “+I” parameter in GTR+I+Γ models—designed to correct for rate heterogeneity and prevent the artificial underestimation of branch lengths. This approach, while computationally convenient, effectively acts as a black box that discards site-specific information, obscuring a primary signature of purifying selection: the absolute persistence of specific nucleotide states at a genomic position across deep time.

The field of genomic functional annotation has since transitioned through three distinct analytical epochs. The first was defined by the mathematical formalization of phylogenetic likelihoods, yielding foundational tools like phyloP (Pollard <u>et al.</u>, 2010) and GERP++ (Davydov <u>et al.</u>, 2010) that provided essential, site-specific maps of purifying selection. While effective for identifying conserved regulatory elements, these methods often encounter a “significance plateau” at invariable sites in deep alignments. In the limit of absolute stasis, standard Likelihood Ratio Tests (LRTs) saturate at a maximal significance ceiling, losing the resolution necessary to distinguish between sites that are invariant due to low evolutionary opportunity and those that are invariant or nearly invariant due to extreme evolutionary constraint. For a detailed technical review of existing methodologies and their respective limitations in the context of site-level conservation, see the Supplementary Material. The second epoch expanded the dimensionality of conservation, utilizing sophisticated null models to identify multi-layered and overlapping regulatory architectures (e.g., FRESCO, Sealfon <u>et al.</u> (2015)). Currently, the field is entering a third epoch dominated by phylogeny-aware deep learning architectures and genomic language models (gLMs) such as GPN-Star (Ye <u>et al.</u>, 2025) and AlphaGenome (Avsec <u>et al.</u>, 2026). These models shift the paradigm from raw nucleotide identity to “semantic matching,” capturing hidden regulatory grammar across hundreds of millions of years. However, these advanced neural networks often act as black boxes, making it difficult to isolate the underlying phylogenetic signal from complex biochemical and contextual features.

We present B-STILL (Bayesian Significance Test of Invariant Low Likelihoods), a framework that bridges these eras by resolving the selective landscape of protein-coding sequence data around the limit of absolute evolutionary stasis. The core intuition of B-STILL is that the significance of an invariant site is conditioned on its cumulative substitution opportunity. By leveraging the site-specific and gene-specific synonymous substitution rate distributions as an internal control for the neutral background substitution rate, B-STILL distinguishes between sites that are invariant because they are evolutionarily cold (low neutral substitution rate) and those that are truly anchored (absolute sequence stasis despite synonymous opportunity). By pooling information across the entire gene to estimate data-specific empirical priors, B-STILL calculates an Empirical Bayes Factor (EBF) that identifies *Evolutionary Stasis Anchors* (ESAs) as positions where strict sequence constraint is statistically anomalous relative to the global selective regime for the entire gene. To identify larger-scale functional footprints, we complement these site-level inferences with a non-parametric Hypergeometric Scan Statistic that detects regional *Stasis Clusters*—contiguous genomic intervals exhibiting a significantly high density of ESAs relative to the gene-wide background.

## Methods

### The B-STILL Framework

B-STILL is built upon the FUBAR (Fast, Unconstrained Bayesian AppRoximation) framework Murrell <u>et al.</u> (2013), which utilizes a Dirichlet process prior to model rate heterogeneity across sites. The underlying evolutionary process is modeled using a standard Muse-Gaut (MG94) codon model modified with a General Reversible (REV) nucleotide substitution matrix Muse and Gaut (1994); Kosakovsky Pond and Frost (2005). The rate of substitution from codon *i* to codon *j* (*q*_*ij*_) is defined by:

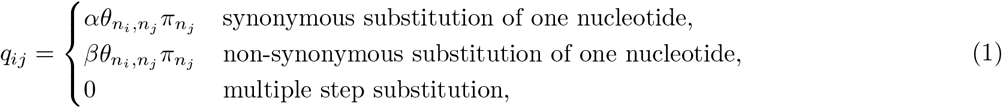

where *α* and *β* represent the site-specific synonymous and non-synonymous substitution rates, *θ* is the nucleotide exchangeability (relative to *θ*_*AG*_ = *θ*_*GA*_ = 1, and *π* is the target nucleotide frequency. The equlibrium frequency over sense codons is obtained using the CF_3×4_ estimator (Kosakovsky Pond <u>et al.</u>, 2010). In the B-STILL implementation, the nucleotide exchangeability parameters (*θ*) and branch lengths are first estimated using a standard GTR model on the entire alignment to provide a robust background fit. These parameters are then held fixed during the subsequent Bayesian inference phase, allowing the model to focus exclusively on site-specific rate variation.

### Fixed-Grid Bayesian Inference (FUBAR)

To achieve computational efficiency, B-STILL follows the FUBAR approach of approximating the continuous rate distribution using a discrete *K* × *K* grid of (*α, β*) points. Let 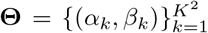 be the set of grid points. The likelihood of the data at site *s* given a specific grid point *k* is *L*_*s*_(*α*_*k*_, *β*_*k*_), which is calculated using the standard Felsenstein’s pruning algorithm. FUBAR avoids the computational burden of traditional MCMC sampling over branch lengths and substitution parameters by leveraging this fixed grid. The gene-wide prior probability of each grid point, 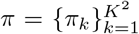, is estimated by pooling information across all *N* sites in the alignment. In B-STILL, we typically employ a 0-th order Variational Bayes (VB0) approximation or a Collapsed Gibbs sampler to estimate the posterior weights *π* that maximize the marginal log-likelihood:

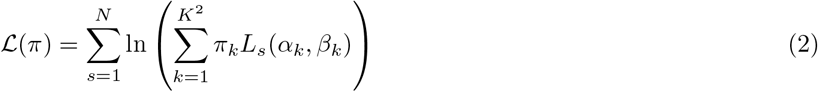

Once the gene-wide prior *π* is established, the posterior probability of site *s* belonging to rate regime *k* is calculated as:

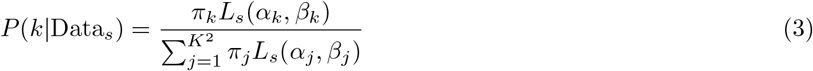

This hierarchical sharing allows B-STILL to calibrate the surprise of invariance at a single site against the background frequency of conservation observed across the entire gene.

Standard selection scans often struggle at the boundary of zero substitutions because linear or log-spaced grids lack the density required to resolve the likelihood surface near the origin. B-STILL overcomes this by implementing a high-resolution quadratic grid in the near-zero regime. For a grid of *K* points per dimension, the rates for the *k*-th point are defined as:

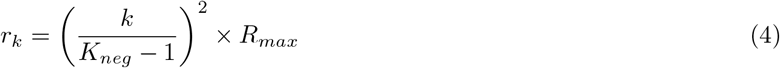

This quadratic clustering provides the statistical sensitivity needed to distinguish between near-zero purifying selection and absolute evolutionary immobilization.

The primary statistical metric in B-STILL is the Empirical Bayes Factor (EBF), which quantifies the surprise of observing a specific selective regime relative to the gene-wide prior.

1. **Exact Invariance (***α* = 0, *β* = 0): Quantifies evidence for absolute sequence constraint.
2. **Synonymous Stasis (***EBF*_*syn*_, *α* = 0): Quantifies evidence for constraint at the nucleotide level, regardless of the non-synonymous rate.
3. **Non-synonymous Invariance (***β* = 0): Quantifies evidence for constraint at the protein level, regardless of the synonymous rate.
4. **Proximal Stasis (***EBF*_*prox*_, *E*[*S*] *< X*): This metric is defined as the posterior probability mass concentrated on grid points (*α, β*) where the total expected number of substitutions along the gene tree is less than or equal to a fixed threshold (e.g., *X* = 0.5, tunable by the user). By explicitly defining a radius of stasis around the origin, B-STILL distinguishes between stochastic invariance and *bona fide* functional constraint.

Unless explicitly defined otherwise, the results and discussion presented herein focus exclusively on Proximal Stasis (*EBF*_*prox*_). The EBF for a state/event *S* is calculated as:

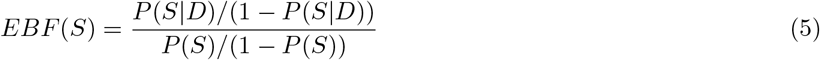

representing the ratio of posterior to prior odds.

### Hypergeometric Scan Statistic for Identifying Stasis Clusters

To identify unusually dense clusters of ESAs within individual coding regions, we implemented a non-parametric clustering algorithm based on an exact Hypergeometric Scan Statistic. Under the null hypothesis (*H*_0_), we assume that ESAs are distributed randomly across the sequence without spatial preference. The gene is treated as a finite population of *L* possible codon positions containing exactly *K* high-confidence ESA sites (e.g., *EBF* ≥ 10).

We define a potential *Stasis Cluster* as any genomic interval [*x*_*i*_, *x*_*j*_] anchored by groups of high-confidence ESAs. For every interval, let *k* be the number of observed ESAs and *d* = *x*_*j*_ − *x*_*i*_ + 1 be the span in codons. The local p-value is calculated using the upper tail of the Hypergeometric distribution:

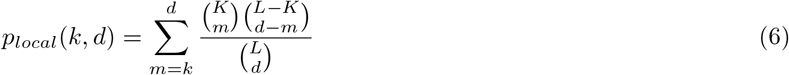

This model accounts for sampling without replacement from a finite population of genomic sites, which is essential when the density of ESAs (*K/L*) is non-negligible. To control the Family-Wise Error Rate (FWER), we performed a permutation test. For each gene, we generated 10,000 permuted datasets by shuffling the positions of the *K* ESAs. The 5th percentile of the null distribution of minimum local p-values was defined as the gene-specific critical significance threshold. Overlapping significant segments were merged into single continuous intervals to demarcate the regional bounds of individual Stasis Clusters.

### Regional Modeling: Comparison with the PHAST Framework

The B-STILL regional scan methodology represents a conceptual departure from established conservation-mapping frameworks such as PHAST (e.g., `phastCons`). While `phastCons` utilizes a Hidden Markov Model (HMM) to identify conserved elements by modeling transitions between “conserved” and “non-conserved” states, it relies on regional rate smoothing to define block boundaries. In contrast, B-STILL identifies Stasis Clusters using a discrete pointprocess model. By anchoring clusters to specific ESAs, B-STILL provides spatial resolution for identifying multilayered features—such as overlapping ORFs or RNA structures—where the requirement for sequence immutability is concentrated in narrow, high-density intervals.

### HIV-1 Structural Mapping and Analysis

To evaluate the protein structural context of ESAs, we projected site-level EBF results onto the crystal structure of the HIV-1 RT heterodimer (PDB: 1RTD; Huang <u>et al.</u> (1998)). Residue-to-residue mapping between the alignment consensus and the PDB sequence was established using global protein alignment (BLOSUM62). Invariant residues were visualized as spheres using the PyMOL Molecular Graphics System (Schrödinger, LLC and DeLano, 2020). We specifically analyzed the spatial clustering of ESAs within the catalytic palm subdomain, encompassing the Asp110–Asp185–Asp186 triad, and the primer-grip motif (residues 227–235).

### Simulation Benchmarks and Power Analysis

To evaluate the statistical properties of the B-STILL framework, we performed power simulations using a mosaic sequence design across 90 distinct evolutionary scenarios, totaling 1,800 replicates. Each simulated alignment was constructed by splicing two selective regimes: a Neutral partition comprising between 90% and 99% of the sites, and a Stasis partition comprising the remaining 1% to 10%. Data were generated using the `simulate` tool in Hy-Phy, with the Neutral partition evolving under gene-specific synonymous and non-synonymous rates (*α, β*) estimated from six empirical datasets: HIV-1 RT (*T* = 6.42, T = tree length in expected substitutions per site), plant RuBisCO (*T* = 10.99, `rbcL`), SARS-CoV-2 Spike (*T* = 0.12), mammalian Beta-globin (*T* = 2.25, `bglobin`), Camelid VHH (*T* = 14.13), and Encephalitis Virus Envelope (*T* = 0.83). For each empirical background, we evaluated performance across five tree-scaling factors (0.1 ×, 0.5×, 1.0×, 2.0×, and 5.0×) and three stasis-injection proportions (0.01, 0.05, and 0.10), generating 10 replicates per scenario. The Stasis partition was simulated by scaling the background tree to model an approximate stasis selective regime, where the upper bound on permitted evolutionary change is lower than the gene-wide average (*E*[*S*] ≤ 0.5). We also included a strict null scenario where 100% of the sites evolved under the background empirical rates without stasis injection to measure the intrinsic False Positive Rate (FPR). Replicates were analyzed using B-STILL with a Proximal Stasis threshold of *EBF* ≥ 10, and performance was assessed by comparing the inferred ESAs to the coordinates of the spliced stasis partition. This design tests the framework’s ability to distinguish functional sequence constraint from stochastic invariance across a spectrum of evolutionary depths and ESA densities.

### Gene Selection for Mammalian Exome Analysis

To evaluate B-STILL’s performance on human-relevant pathology, we targeted a definitive set of genes harboring well-documented, disease-causing synonymous variants. Candidate genes were selected based on an exhaustive review of the clinical genetics literature and databases (e.g., ClinVar, HGMD), focusing on positions where synonymous single nucleotide variants (sSNVs) have been definitively established as pathogenic drivers through *in vitro* or *in vivo* validation.

The selection process prioritized four primary molecular mechanisms of synonymous pathogenicity: (1) disruption of pre-mRNA splicing networks (e.g., `MSH2`, `TP53`, `BRCA1`), (2) perturbation of co-translational protein folding and translation kinetics (e.g., `CFTR`, `F9`), (3) alteration of mRNA thermodynamic stability (e.g., `DRD2`), and (4) destruction or creation of microRNA (miRNA) binding sites (e.g., `IRGM`). This yielded a validation panel of 38 high-priority genes representing diverse functional classes, including tumor suppressors, coagulation factors, and metabolic enzymes.

### Comparative Benchmarking against phyloP

To isolate the methodological advantage of B-STILL’s codon-aware framework, we conducted a head-to-head comparison against phyloP Pollard <u>et al.</u> (2010) using a validation panel of 38 representative mammalian genes. This panel was selected to encompass a spectrum of selective regimes, ranging from ultra-conserved housekeeping genes (e.g., `CALM1`) to fast-evolving antiviral and surface proteins (e.g., `APOBEC3D`, `MUC1`), as well as established clinical targets (`TP53`, `BRCA1`, `CFTR`). To ensure a comparison, phyloP was executed on the identical 120-species mammalian alignments and neutral phylogenetic backgrounds used for the B-STILL analysis. Specifically, for each gene, we first estimated a synonymous-aware neutral tree using the FitMG94 framework (`FitMG94.bf`) Kosakovsky Pond and Frost (2005), extracting the expected synonymous substitutions per nucleotide site across the entire phylogeny. This tree was utilized as a fixed constraint in phyloFit Siepel <u>et al.</u> (2005) to estimate gene-specific General Time Reversible (GTR) substitution parameters and equilibrium frequencies (`--estimate-freqs`) while holding branch lengths constant (`--no-opt branches`). Site-specific conservation and acceleration scores were subsequently calculated using phyloP with the Likelihood Ratio Test (`--method LRT`) in `CONACC` mode.

### Dark Proteome Analysis

To evaluate the utility of B-STILL for the functional annotation of uncharacterized proteins, we performed a proteome-wide scan of the “dark proteome”—defined here as the subset of the human genome consisting of uncharacterized open reading frames (ORFs) and poorly characterized gene families. Our analysis targeted 815 genes from the 120-species mammalian alignment, including the `C*orf` (284 genes), `FAM` (228 genes), `TMEM` (272 genes), and `KIAA` (31 genes) nomenclature groups. These genes were prioritized based on the absence of experimentally validated biochemical functions or detailed structural characterization in major databases (e.g., UniProt, PDB). For each gene, we identified ESAs and spatial clusters (*EBF* ≥ 100, increased stringency). To investigate the structural implications of these ESAs, site-specific Bayesian factors were mapped onto AlphaFold-predicted 3D structures. The spatial distribution of selective constraint was visualized by substituting the B-factor column of the corresponding PDB files with log_10_(*EBF*_*prox*_). This approach allowed for the identification of physically clustered ESAs in three-dimensional space, providing a data-driven strategy for prioritizing structural hubs and potential interaction interfaces in proteins that have otherwise eluded experimental characterization.

### Validation against Human Population Variation

We validated the biological relevance of mammalian ESAs by cross-referencing our phylogenetic inferences with human population-level variation from the gnomAD database (v4.1.1; Chen <u>et al.</u> (2024)). For each analyzed mammalian gene (a subset of *N* = 68), we mapped human protein coordinates to the 120-species alignment consensus (Hecker and Hiller, 2020) using global protein alignment (BLOSUM62) and the HGVSp nomenclature Hart <u>et al.</u> (2024). We retrieved non-synonymous and synonymous variants and quantified the relationship between B-STILL EBFs and human allele frequencies using the Spearman rank correlation coefficient. To assess clinical utility, we performed a receiver operating characteristic (ROC) analysis using pathogenic and benign variants from the ClinVar database Landrum <u>et al.</u> (2018) for a further subset of these genes chosen for known clinical significance (*N* = 18). For this analysis, ClinVar clinical significance was used as the binary classification target (Pathogenic and Likely Pathogenic variants were assigned a value of 1; Benign and Likely Benign variants were assigned a value of 0). We utilized B-STILL EBFs as continuous predictors to assess their ability to distinguish between clinically confirmed deleterious substitutions and neutral variation. Specifically, we calculated the Area Under the Curve (AUROC) using *EBF*_*prox*_ for non-synonymous variants and *EBF*_*syn*_ for synonymous variants. Only high-confidence variants (reviewed by an expert panel or with multiple submitters and no conflicting interpretations) were included in the validation panel to ensure a rigorous clinical ground truth.

### REVEL Correlation and Clinical Benchmarking

To further evaluate the relationship between B-STILL EBFs and established clinical pathogenicity predictors, we performed a genome-scale comparison with the REVEL (Rare Exome Variant Ensemble Learner) ensemble method Ioannidis <u>et al.</u> (2016). We selected a validation panel of 85 genes by identifying the top 100 fastest-evolving genes in our mammalian dataset (ranked by total tree length) and filtering for those with available non-synonymous variant scores in the MyVariant.info database Louter <u>et al.</u> (2016). This strategy targeted high-divergence gene families—including Zinc Fingers (ZNF), Olfactory Receptors (OR), and Taste Receptors (TAS2R)—to evaluate B-STILL’s performance in selective regimes with high mutational opportunity. The resulting dataset encompassed 34,206 individual non-synonymous variant positions. REVEL scores and protein-level coordinates were retrieved programmatically using the `MyVariant.info` REST API. To ensure precise coordinate integration, human protein positions were mapped to the 120-mammal MSA using translated `hg38` sequences, accounting for local gaps and indels.

### Analysis of Overlapping Reading Frames

To evaluate the performance of B-STILL in detecting multi-layered functional constraints, we analyzed 111 viral protein-coding alignments from the FRESCO dataset (Sealfon <u>et al.</u>, 2015). For each alignment, we computed sitespecific EBFs for proximal stasis (*α* ≈ 0, *β* ≈ 0). To validate these signals, we curated reference genomes from the NCBI Virus RefSeq database (e.g., NC 004102 for Hepatitis C, NC 011505 for Rotavirus A). Annotated overlapping reading frames were defined as any two CDS features sharing genomic coordinates but translated in different frames or on opposite strands. Nucleotide-aware projection (BLOSUM62) was used to map RefSeq coordinates to alignment indices, accounting for gaps and frame-shifts.

### Implementation and Availability

B-STILL is implemented as a standard analysis module within the HyPhy software package (version 2.5.95 or later) using the HyPhy Batch Language (HBL). The implementation leverages HyPhy’s high-performance likelihood engine and supports both traditional grid-based posterior estimation (via a Collapsed Gibbs sampler) and rapid 0-th order Variational Bayes inference (VB0). The framework introduces a configurable `radius-threshold` parameter, which corresponds to the *X* value described in the methodology. This parameter allows users to define proximal constraint in terms of total evolutionary surprise across the tree. For example, setting *X* = 0.5 effectively classifies any rate regime expected to produce fewer than one substitution across the entire phylogeny as belonging to the proximal invariant state. This approach is critical for distinguishing between sites that are “historically invariant” (zero substitutions observed) and those that are “effectively invariant” (substitution rates significantly below the gene-induced expectation even if rare substitutions are present). Resulting EBFs and model parameters are exported in a structured JSON format, compatible with the HyPhy vision ecosystem for interactive visualization. Significant regional Stasis Cluster signals were identified using a standalone Python tool, `infer_stasis_clusters.py`. This tool implements the exact Hypergeometric Scan Statistic described above, performing 10,000 permutations per gene to determine gene-specific critical p-value thresholds for family-wise error rate control. The source code for B-STILL and the Hypergeometric Scan Statistic is available under the MIT license at https://github.com/veg/hyphy and https://github.com/veg/b-still. B-STILL is also available through a user-friendly web interface on Datamonkey v3 (v3.datamonkey.org) and as a tool within the Galaxy Project ecosystem.

## Results

### HIV-1 RT Case Study

The fundamental strength of the B-STILL framework lies in its ability to resolve site-specific selective regimes by leveraging both gene-wide information and local mutational/substitution opportunity. This process is illustrated by comparing the global rate distribution (the prior) with the inferred posterior distribution for a site under extreme evolutionary stasis (Figure 1). While the gene-wide prior for HIV-1 Reverse Transcriptase reflects a range of synonymous and non-synonymous rates (Figure 1A), B-STILL identifies sites where the local data effectively pulls the probability mass toward the (0, 0) origin (Figure 1B). Our results reveal that not all invariant sites are equal. The EBF effectively measures the surprise of observed stasis relative to a codon-aware gene-wide baseline. A codon with high synonymous redundancy, such as Serine (`TC*`), faces a significantly higher expected rate of synonymous drift than a codon with low redundancy, such as Tyrosine (`TA[T/C]`). Under a neutral model, the probability of observing zero substitutions at a site with high synonymous opportunity is relatively smaller compared to a site where substitution targets are restricted by the genetic code. Consequently, B-STILL yields a substantially higher EBF for invariance at highly redundant codons, as their stasis represents a more substantial deviation from the gene-wide expectation.

**Figure 1:**
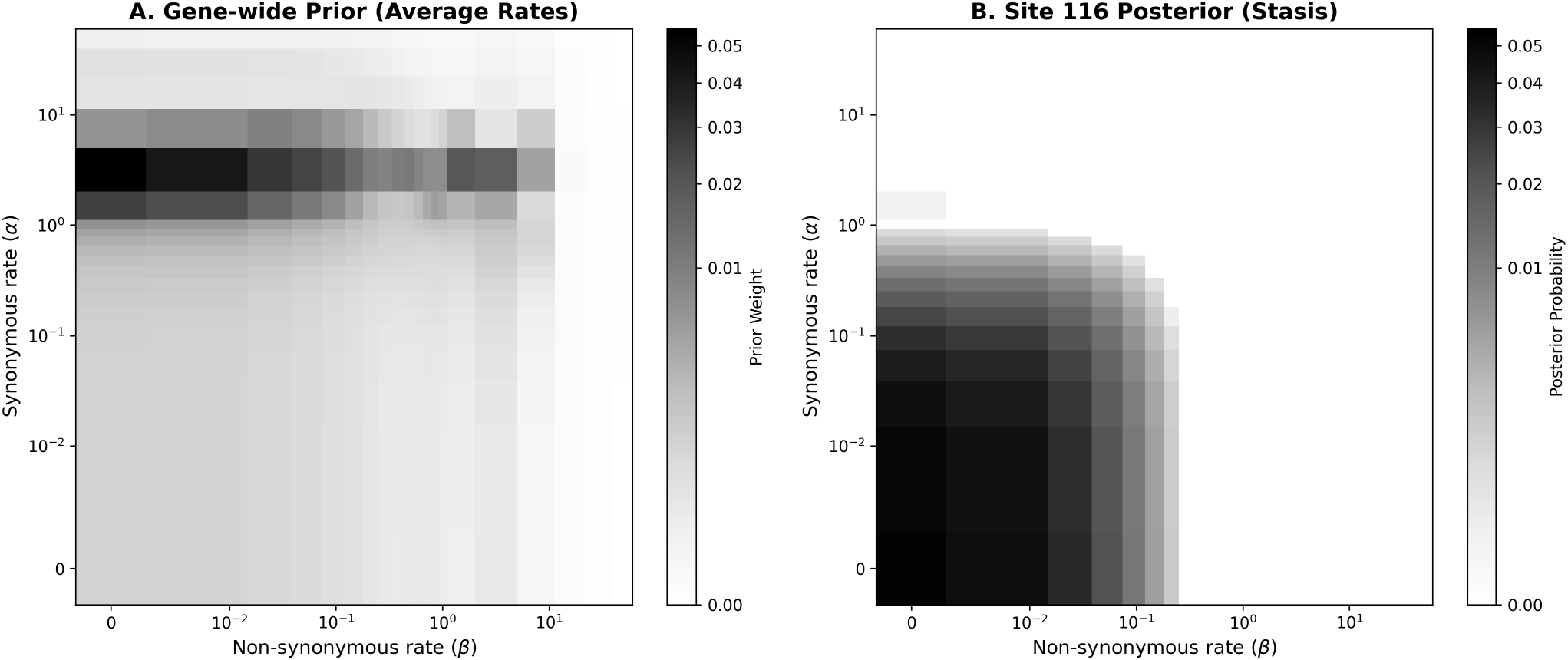
Bayesian resolution of evolutionary stasis in HIV-1 Reverse Transcriptase. (A) The gene-wide prior distribution of synonymous (*α*) and non-synonymous (*β*) substitution rates, representing the average selective landscape across the entire gene. (B) The posterior distribution for site 116, showing the concentration of probability mass at the (0, 0) origin. This transition from a broad prior to a peaked posterior at the origin is the hallmark signature of an Evolutionary Stasis Anchor.

Furthermore, the method accounts for the underlying nucleotide substitution matrix. If the inferred GTR parameters indicate a high transition/transversion ratio (*κ*), a codon whose synonymous targets require a transition is expected to vary more than one requiring a transversion. For example, comparing site 117 and site 232 in RT illustrates this resolution. Site 117 (Serine, `TCA`) resides in a 4-fold degenerate block with multiple high-probability transition paths to synonymy. Despite the presence of mixed bases (`TCR` and `TMA`) which represent synonymous ambiguities, the total lack of non-synonymous change is statistically significant (*EBF*_117_ = 1990.84). In contrast, site 232 (Tyrosine, `TAT`) has a much narrower synonymous neighborhood (*S* = 1). While it also exhibits an ambiguity (`TAY`), its stasis is less surprising under the hierarchical prior, resulting in a lower, albeit still significant, EBF (107.39).

By explicitly calculating the synonymous and non-synonymous stencils, we can partition the total constraint into its respective components, revealing sites that are under intense non-synonymous purifying selection despite being synonymous neutral. Projection of invariant sites onto the RT crystal structure further corroborates their functional relevance. Invariant residues cluster in catalytically and structurally critical regions, including the palm subdomain and the primer-grip motif (Figure 2). This spatial enrichment provides independent structural validation that B-STILL proximal-constraint calls correspond to functional immobilization.

**Figure 2:**
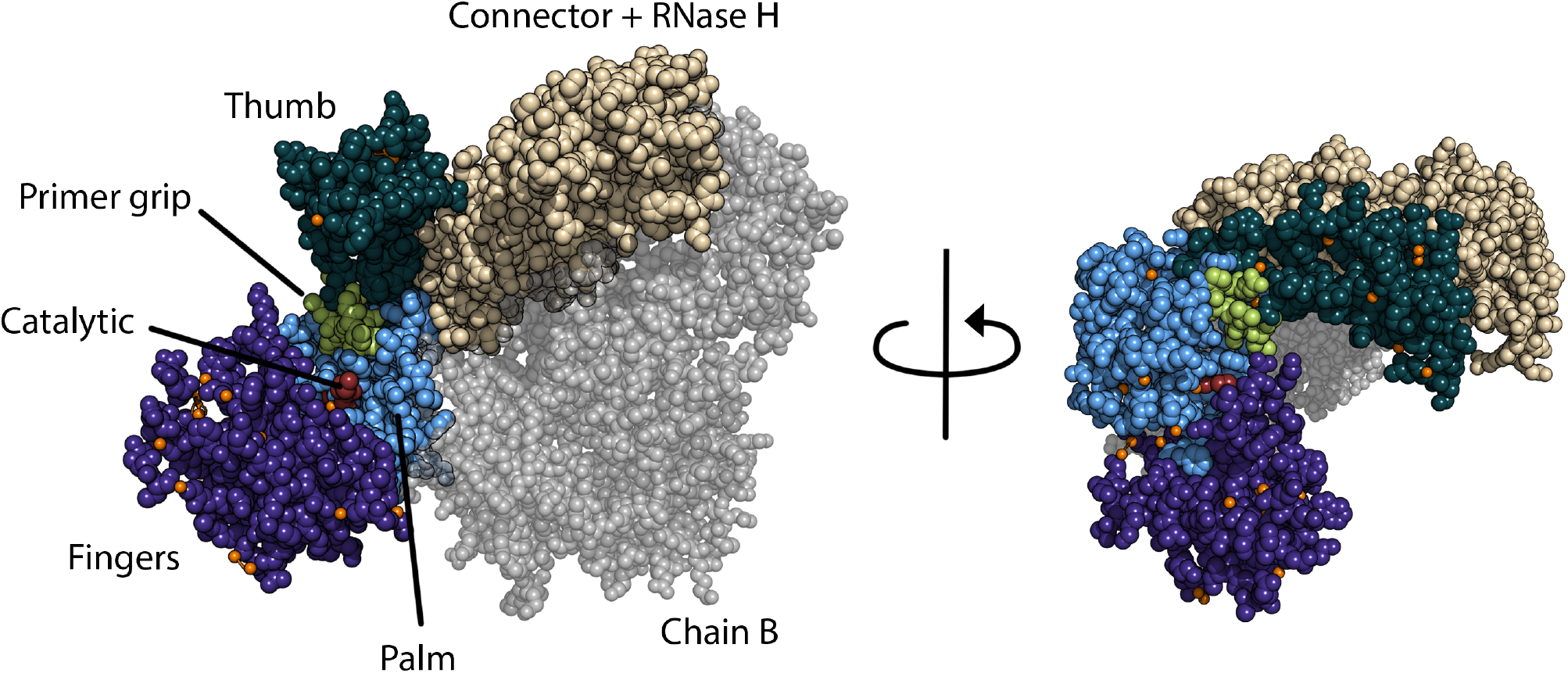
Structural mapping of invariant sites in HIV-1 Reverse Transcriptase (PDB: 1RTD) with a global view of B-STILL inferred invariant residues (red spheres) projected onto the RT heterodimer.

As an example of extrinsic validation, we reasoned that sites inferred as highly constrained by B-STILL from this training dataset would exhibit lower variability in a large population sample. Indeed, examining a population database of over 175,000 HIV-1 sequences (Stanford HIVDB; Rhee <u>et al.</u> (2003)), we found that evolutionary stasis is a predictor of human population-level variability: codons with higher B-STILL EBFs had lower frequencies of amino-acid mutations (*ρ* = − 0.3271, *p <* 10^−9^). Significant proximal ESAs identified in the RT alignment (i.e. those with associated EBFs ¿10) are summarized in Table 1.

**Table 1:** Sites in HIV-1 RT inferred to be under significant proximal constraint (*EBF* ≥ 10). *α* and *β* represent the mean posterior synonymous and non-synonymous rates, respectively.

| Site | $\alpha$ | $\beta$ | $P[\text{prox}]$ | $EBF[\text{prox}]$ | Codon Composition |
| --- | --- | --- | --- | --- | --- |
| 4 | 0.701 | 0.138 | 0.6586 | 12.47 | CCT (162), CCC (4), YCT (1), ... |
| 5 | 0.357 | 0.048 | 0.8946 | 54.89 | ATT (167), RTT (1) |
| 7 | 0.287 | 0.040 | 0.9338 | 91.18 | ACT (183), RCT (1) |
| 10 | 0.676 | 0.039 | 0.7715 | 21.83 | GTA (182), GTG (2) |
| 14 | 0.244 | 0.036 | 0.9540 | 134.08 | CCA (184) |
| 17 | 0.515 | 0.039 | 0.8221 | 29.87 | GAT (200) |
| 25 | 0.590 | 0.127 | 0.7520 | 19.61 | CCA (200), GCA (2), CCR (1), ... |
| 27 | 0.657 | 0.034 | 0.7896 | 24.26 | ACA (201), ACG (2), WCA (1), ... |
| 30 | 0.828 | 0.041 | 0.6708 | 13.17 | AAA (201), AAR (3), AAG (1) |
| 31 | 0.922 | 0.030 | 0.6422 | 11.60 | ATA (204), WTA (2) |
| 33 | 0.233 | 0.036 | 0.9580 | 147.49 | GCA (206) |
| 51 | 0.765 | 0.017 | 0.8497 | 36.53 | GGG (460), GGA (14), GGT (2) |
| 52 | 0.671 | 0.017 | 0.8269 | 30.89 | CCT (472), CCC (2), CCW (2) |
| 59 | 0.775 | 0.017 | 0.7660 | 21.16 | CCA (469), CCG (4), CCR (3) |
| 105 | 0.579 | 0.025 | 0.8871 | 50.81 | TCA (471), TCG (3), TCR (2) |
| 113 | 0.729 | 0.020 | 0.7516 | 19.56 | GAT (465), RAT (6), GAC (2), ... |
| 117 | 0.102 | 0.028 | 0.9968 | 1990.84 | TCA (474), TCR (1), TMA (1) |
| 129 | 0.731 | 0.071 | 0.7673 | 21.32 | GCA (464), GCC (4), GCT (3), ... |
| 137 | 0.726 | 0.021 | 0.7524 | 19.65 | AAT (471), AAC (2), ART (1), ... |
| 140 | 0.586 | 0.018 | 0.8863 | 50.39 | CCA (470), CCM (2), CCC (2), ... |
| 156 | 0.923 | 0.022 | 0.6183 | 10.47 | TCA (465), TCR (4), TCG (4), ... |
| 185 | 0.270 | 0.022 | 0.9470 | 115.56 | GAT (476) |
| 189 | 0.096 | 0.158 | 0.9796 | 310.99 | GTA (470), ATA (2), GTR (2), ... |
| 209 | 0.945 | 0.031 | 0.6617 | 12.64 | CTG (387), CTA (37), TTG (25), ... |
| 217 | 0.106 | 0.020 | 0.9968 | 1986.41 | CCA (464), CCW (5), CCM (1) |
| 222 | 0.845 | 0.034 | 0.7362 | 18.04 | CAG (453), CAA (13), CAR (3) |
| 232 | 0.274 | 0.030 | 0.9432 | 107.39 | TAT (467), TAY (2) |
| 240 | 0.934 | 0.018 | 0.6106 | 10.14 | ACA (453), ACG (4), ACR (2), ... |
| 247 | 0.636 | 0.019 | 0.8519 | 37.19 | CCA (428), CCT (2), SCA (1), ... |
| 254 | 0.514 | 0.052 | 0.8658 | 41.72 | GTC (146), GTT (2) |
| 255 | 0.627 | 0.051 | 0.7587 | 20.33 | AAT (148) |
| 290 | 0.903 | 0.046 | 0.6222 | 10.65 | ACA (117), ACG (2) |
| 299 | 0.447 | 0.061 | 0.8379 | 33.42 | GCA (89), GSA (2), GCR (2) |
| 300 | 0.707 | 0.071 | 0.7107 | 15.88 | GAG (86), GAA (5), GRG (1) |
| 310 | 0.454 | 0.219 | 0.6793 | 13.70 | CTA (35), YTA (1) |

### Simulation Benchmarks: Specificity and Depth-Dependent Power

Benchmarks across 1,800 nucleotide sequence datasets that were simulated using the phylogenies of six representative genes under three different scenarios, confirm that B-STILL is conservative, maintaining a Null False Positive Rate (FPR) ≪ 1% across all datasets (Figure 3C)(Table 2).

**Table 2:** Summary of B-STILL performance across 900 simulation replicates (50 per Gene/Scenario combination). Scenario A represents the neutral null model; Scenario B evaluates extreme purifying selection (*ω* = 0.1) at all sites; Scenario C tests sensitivity under reduced evolutionary depth (0.5× tree scaling).

| Gene | Scenario | Mean Sig | Std Dev | Max Sig |
| --- | --- | --- | --- | --- |
| ENCenv | A | 0.46 | 0.95 | 4 |
| ENCenv | B | 1.32 | 1.45 | 7 |
| ENCenv | C | 0.00 | 0.00 | 0 |
| HIV_RT | A | 0.60 | 0.90 | 5 |
| HIV_RT | B | 1.36 | 1.10 | 5 |
| HIV_RT | C | 3.56 | 2.26 | 10 |
| SARS-CoV-2-spike | A | 0.00 | 0.00 | 0 |
| SARS-CoV-2-spike | B | 0.00 | 0.00 | 0 |
| SARS-CoV-2-spike | C | 0.00 | 0.00 | 0 |
| bglobin | A | 0.30 | 0.51 | 2 |
| bglobin | B | 0.02 | 0.14 | 1 |
| bglobin | C | 0.12 | 0.33 | 1 |
| camelid | A | 0.00 | 0.00 | 0 |
| camelid | B | 0.06 | 0.24 | 1 |
| camelid | C | 0.00 | 0.00 | 0 |
| rbcL | A | 0.00 | 0.00 | 0 |
| rbcL | B | 0.12 | 0.33 | 1 |
| rbcL | C | 0.48 | 0.68 | 2 |

**Figure 3:**
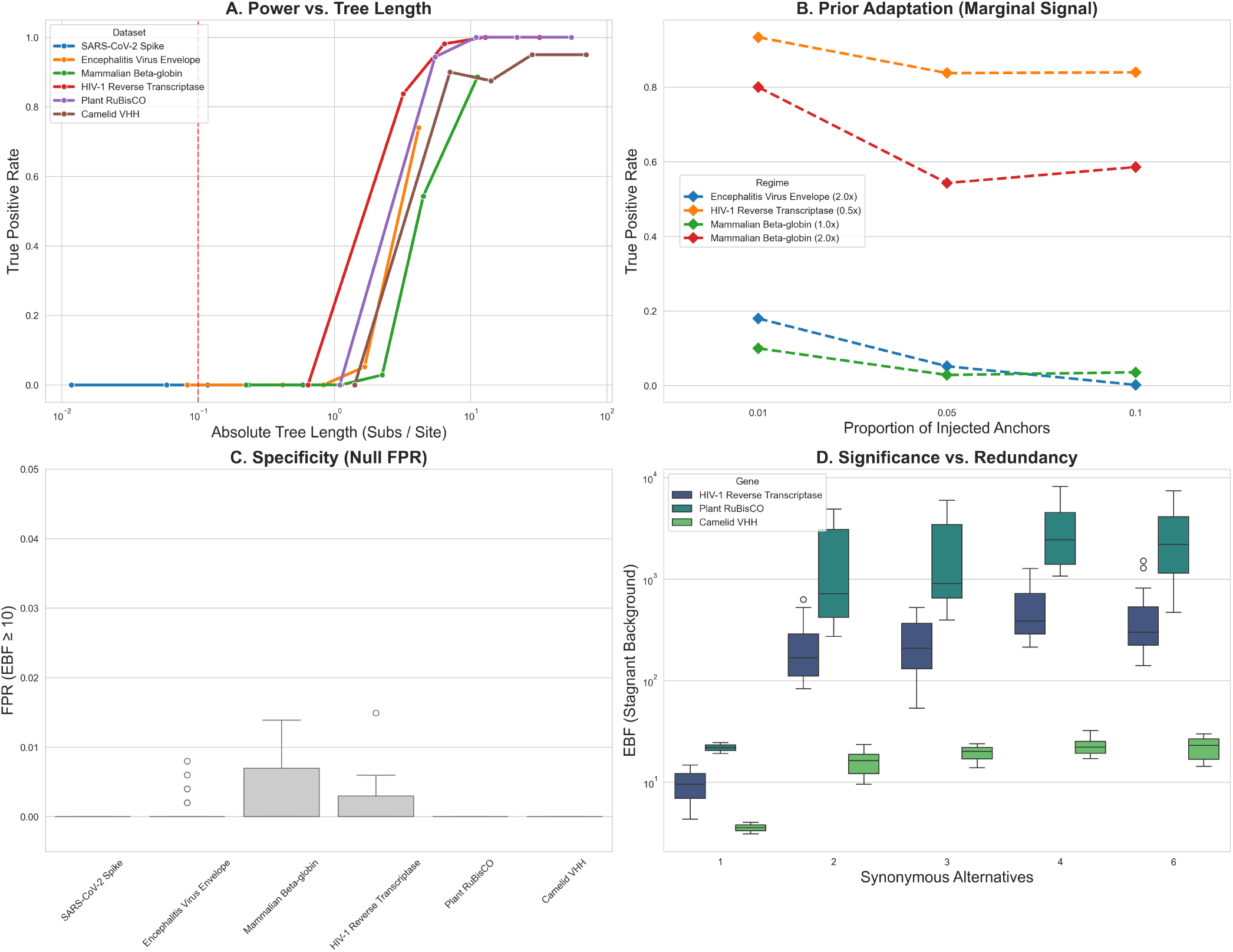
B-STILL performance summary across 1,800 simulated datasets. (A) Detection power (TPR) as a function of absolute tree length (at 5% injected ESA density). Sensitivity scales log-linearly with total substitution count, reaching nearsaturation in deep phylogenies (*T >* 5). (B) Power vs. Anchor Density. Power decreases as the proportion of ESAs increases in marginal signal regimes (e.g., Encephalitis Virus Envelope and Mammalian Beta-globin), reflecting the adaptation of the Bayesian prior to a more evolutionarily stagnant background. (C) FPR control remains robustly below 1% across all datasets. (D) Significance vs. Codon Redundancy. Absolute stasis is statistically more significant at redundant codons.

In the strict null simulation scenario (100% background sites), the framework identified near-zero false positives, confirming that the Bayesian prior successfully adapts to the background mutational process. In datasets simulated using the extremely shallow SARS-CoV-2 Spike phylogeny (*T* = 0.12), B-STILL correctly identified zero ESAs (proximal EBF for *X* = 0.5 expected substitutions per site), confirming that the absence of variation in young lineages is correctly de-weighted as statistically uninformative (Figure 3A). These benchmarks reveal that B-STILL successfully utilizes the redundancy of the genetic code as an internal control for mutational opportunity. We observed that absolute stasis is statistically more significant at redundant codons (e.g., 4-fold Proline) than at low-redundancy ones (e.g., Methionine), reflecting the greater deviation from expectation when substitutions are absent despite opportunities for synonymous variation (Figure 3D).

The power to detect ESAs is driven by evolutionary contrast. We evaluated performance across datasets that were simulated using six empirical phylogenies representing a range of different tree lengths (*T*): HIV-1 RT (*T* = 6.42), RuBisCO (*T* = 10.99), SARS-CoV-2 Spike (*T* = 0.12), mammalian Beta-globin (*T* = 2.25), Camelid VHH (*T* = 14.13), and Encephalitis Virus Envelope (*T* = 0.83). For deep phylogenies (HIV-1 RT, RuBisCO, Camelid VHH), B-STILL identifies ESAs with near-perfect sensitivity (TPR ≈ 100%) while maintaining exceptional specificity (Figure 3A). For phylogenies of moderate depth, such as Beta-globin and Encephalitis Envelope, TPR scales loglinearly with tree length, reaching *>* 70% when background variation is scaled to 5 × the empirical tree depth.

Sensitivity benchmarks reveal that self-calibration through prior adaptation is most pronounced in the marginal signal regime (Figure 3B). In datasets simulated using phylogenies of moderate depth (Encephalitis Virus Envelope and Mammalian Beta-globin), we observe a consistent decrease in detection power as the proportion of ESAs increases. This phenomenon demonstrates the hierarchical prior’s role in governing conservatism: when stasis is rare, any invariant site is highly statistically surprising, but as the gene-wide background becomes more stagnant, stasis at any given site becomes less surprising and the threshold for significance naturally increases. This effect disappears in datasets simulated using deep phylogenies (HIV-1 RT and RuBisCO), where the signal of functional immobilization is strong enough to saturate detection power regardless of the density of ESAs.

Finally, as suggested by the HIV-1 RT analysis, synonymous codon redundancy partially informs the detection of ESAs. For sites simulated under stasis, levels of statistical surprise compared to the background of the gene were higher for codons with higher synonymous redundancy (Figure 3D). This finding confirms the intuition that when a highly synonymously redundant site harbors an unexpectedly small number of substitutions, it is more remarkable than when the same behavior is seen at a less redundant site.

### Global Scan of the Mammalian Exome

We applied B-STILL to 19,117 protein-coding genes from the 120-species mammalian alignment, totaling 10,990,498 codon sites (Hecker and Hiller, 2020). This large-scale scan identifies sites where stasis is anomalous relative to the gene-wide selective background, providing a high-resolution map of ESAs across the proteome (Figure 4).

**Figure 4:**
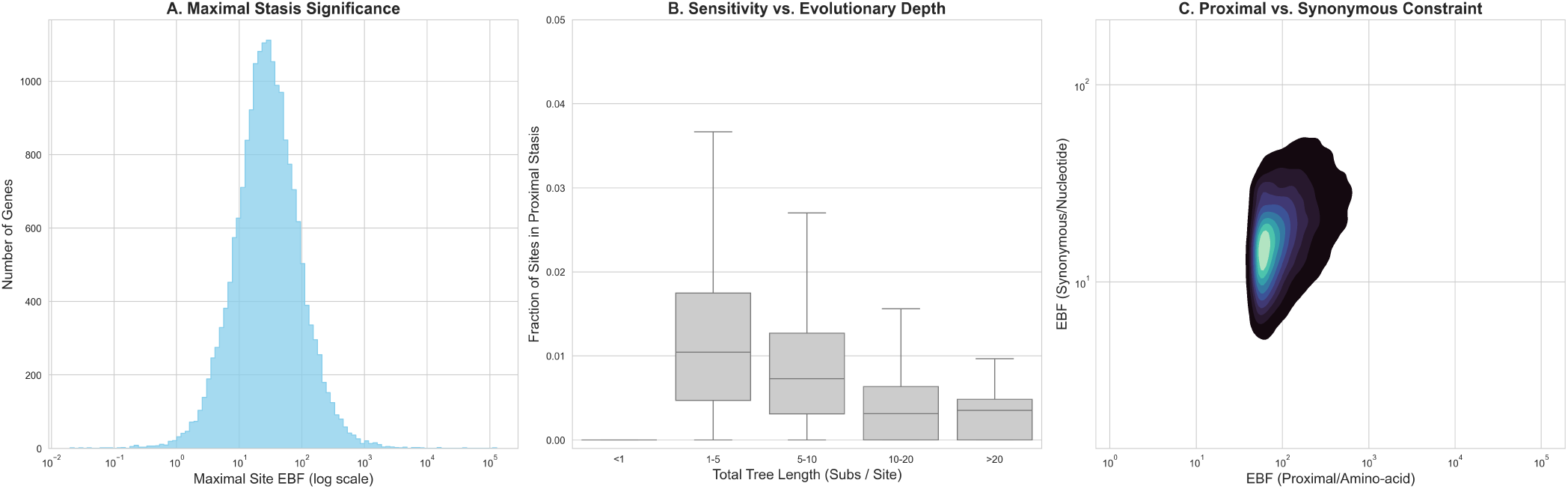
Patterns of evolutionary stasis across 19,117 mammalian genes. (A) Distribution of maximal stasis significance. Isolated ESAs in variable genes reach EBF values (*>* 10^6^). (B) Detection Sensitivity vs. Depth. Signals of evolutionary stasis are negligible in shallow phylogenies (*T <* 1) and peak in phylogeneies of moderate depth (*T* = 1–5). (C) Selective Hierarchy. Absolute sequence constraint (*α, β* ≈ 0) provides a more powerful signal of deviation from expectation than synonymous-only constraint.

The prevalence of ESAs exhibits a characteristic non-monotonic relationship with evolutionary depth. Our results demonstrate that detection sensitivity is negligible in shallow phylogenies (tree length *T <* 1 substitution per site, Figure 4B), where evolutionary stasis is statistically expected. The detectability of ESAs peaks in phylogenies of moderate depth (*T* between 1–5 substitutions per site, Median Proportion of static sites = 1.04%), providing the optimal contrast for resolving selective constraint across the proteome. Notably, the proportion of identified ESAs declines progressively as evolutionary depth increases, falling to 0.35% in the deepest phylogenies (*T >* 20). However, the intensity of stasis significance (maximal site EBF) scales positively with tree length, with the mean maximal EBF rising from 2.44 in shallow phylogenies to over 4, 500 in the deepest phylogenies (not shown). This trend confirms that while the requirement for absolute sequence immutability through deep time is rare, the observation of such stasis in high-divergence alignments provides a powerful signal of functional evolutionary constraint.

Across the mammalian exome, 151,146 sites (1.38%) exhibit significant evidence of proximal stasis (*EBF*_*prox*_ ≥ 10), while only 74,080 sites (0.67%) achieve the same threshold for synonymous-only constraint (*EBF*_*syn*_ ≥ 10). This characteristic hierarchy in significance reflects the framework’s focus on statistical surprise. Because the prior expectation for absolute sequence constraint is an order of magnitude lower than for synonymous stasis, the observation of a perfectly constant site provides a significantly larger signal of deviation from expectation (Figure 4C).

The prevalence of detectable ESAs is non-uniformly distributed across the genome. While the average gene contains approximately 35 significant proximal ESAs, a subset of genes—often those involved in core developmental processes, RNA regulation, or structural stabilization—contain “islands” of extreme conservation where every site is effectively immobilized. As predicted by B-STILL’s hierarchical prior, the statistical “surprise” of a given invariant site is strongly dependent on the background mutation rate of the gene. In fast-evolving genes such as olfactory receptors or mucins, isolated invariant sites achieve astronomical EBF values (*>* 10^6^), representing ESAs that maintain critical structural or functional roles despite massive diversifying pressure on the rest of the protein (e.g., `OR4N5` Site 169, EBF = 8.03 × 10^8^). Conversely, in ultra-conserved genes like Histones or Calmodulin, the high prior expectation of invariance de-weights individual sites, appropriately identifying them as part of a globally constrained architecture rather than specific anomalous ESAs. These findings establish B-STILL as a tool for distinguishing between uninformative sequence conservation and the operational core of protein function.

### Validation against Human Variation and Clinical Ground Truth

We cross-referenced the sites found by B-STILL in the scan of a panel of 68 mammalian genes (see methods) against both degrees of variation at those sites in the human population (using gnomAD v4.1.1), and evidence of mutations at the sites having pathological consequences in humans (using ClinVar and REVEL). Our analysis revealed statistically significant negative correlations between mammalian EBFs and human allele frequencies for the majority of genes, with the strongest predictive power observed for genes like `VHL` and `F9` (Table 3).

**Table 3:**
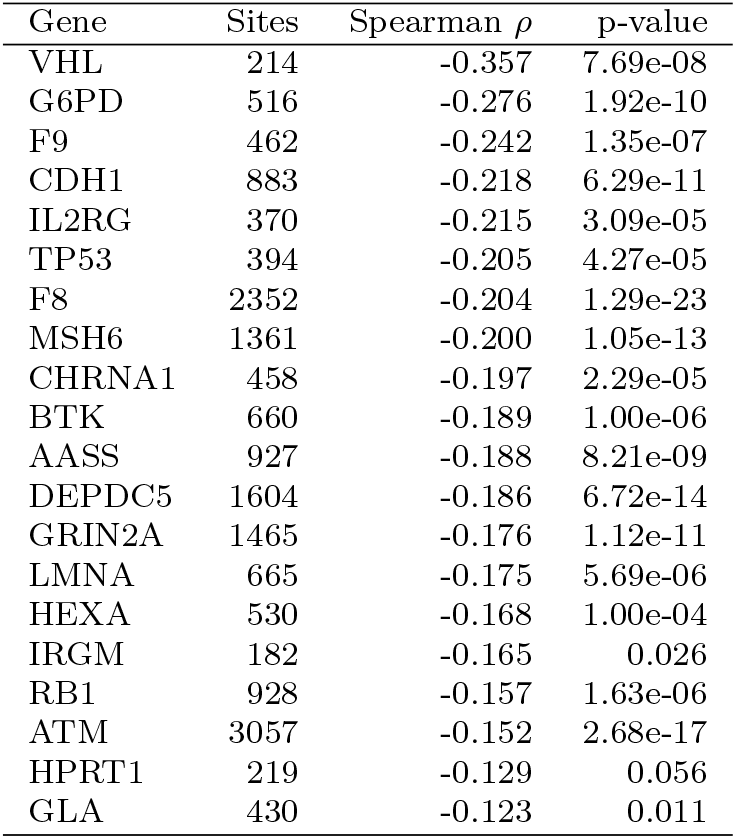
Validation of B-STILL Proximal EBFs against human population variation (gnomAD v4.1). *ρ* represents the Spearman correlation between site-level EBF and the sum of allele frequencies at that site. Top 20 genes by correlation magnitude are shown.

B-STILL EBFs demonstrate strong predictive power with respect to predicting clinical pathogenicity (Figure 5) in a subset of 18 genes selected for harboring known pathogenic variants. For non-synonymous variants, the B-STILL framework yields an aggregate AUROC of 0.65 (mean per-gene AUROC = 0.71). More notably, B-STILL uniquely identifies evidence of clinically relevant synonymous variants, achieving an aggregate AUROC of 0.88 (mean pergene AUROC = 0.89) using *EBF*_*syn*_, identifying evolutionary constraints on mutations in regulatory sequences that are embedded within coding regions: constraints that are invisible to AI models that focus exclusively on protein sequences.

**Figure 5:**
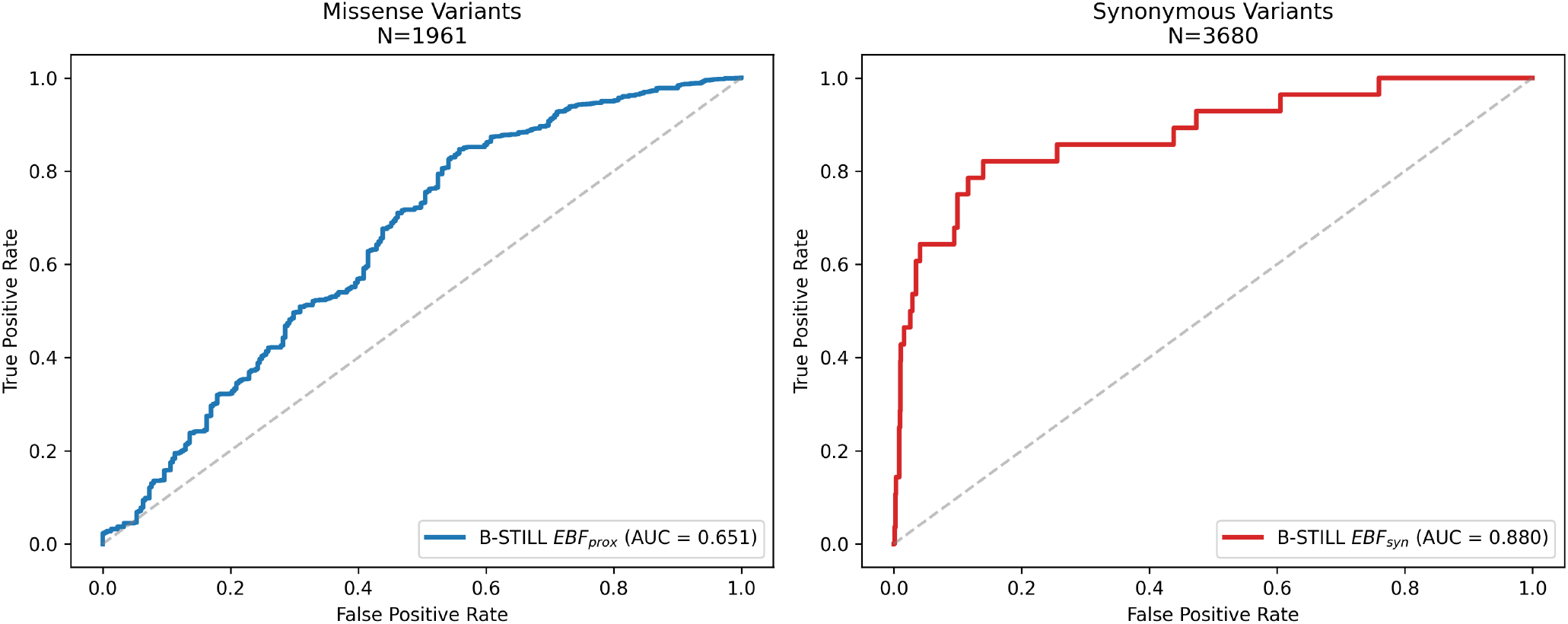
Clinical validation of B-STILL Empirical Bayes Factors using pathogenic and benign variants from ClinVar. (Left) Aggregate ROC curve for non-synonymous variants (AUROC = 0.65), using *EBF*_*prox*_ as the predictor. (Right) Aggregate ROC curve for synonymous variants (AUROC = 0.88) using *EBF*_*syn*_. The True Positive Rate (TPR) represents the proportion of clinically confirmed pathogenic variants correctly identified as Evolutionary Stasis Anchors, while the False Positive Rate (FPR) denotes the proportion of confirmed benign variants incorrectly flagged. These results demonstrate that Evolutionary Stasis Anchors are a powerful and specific predictor of clinical pathogenicity, particularly for synonymous regulatory variation.

For the panel of *N* = 85 reference genes, B-STILL EBFs show a strong positive correlation with REVEL pathogenicity scores (*ρ* = 0.40, *p <* 10^−300^, Figure 6). This concordance is noteworthy given that ESAs represent only the most extreme subset of the proteome: a conservative marker that captures only a fraction of the total landscape of clinical pathology. Furthermore, while REVEL is explicitly designed to score non-synonymous substitutions and cannot therefore evaluate non-protein-altering variation, B-STILL leverages a single phylogenetic signal of absolute sequence constraint that applies to both non-synonymous and synonymous positions. Despite its limited scope relative to REVEL’s 13-feature ensemble, B-STILL correctly identifies a shared functional core of genome sites where extreme evolutionary stasis and clinical consensus align, while also highlighting “Hidden Anchors” where intense purifying selection precedes or complements clinical consensus. For instance, at site 351 of `FGFR3` where mutations are known to cause hypochondroplasia Yao <u>et al.</u> (2019), B-STILL identifies a high-confidence ESA (*EBF* = 215.9) while the REVEL score remains low (0.15). Additionally, B-STILL identifies critical clinical constraint at sites where substitutions are strictly synonymous in humans, such as `TP53` site 125 (*EBF*_*prox*_ = 110) and `CFTR` site 1239 (*EBF*_*syn*_ = 109), which map to documented regulatory pathogenic variants Pinto <u>et al.</u> (2022)Molinski <u>et al.</u> (2014).

**Figure 6:**
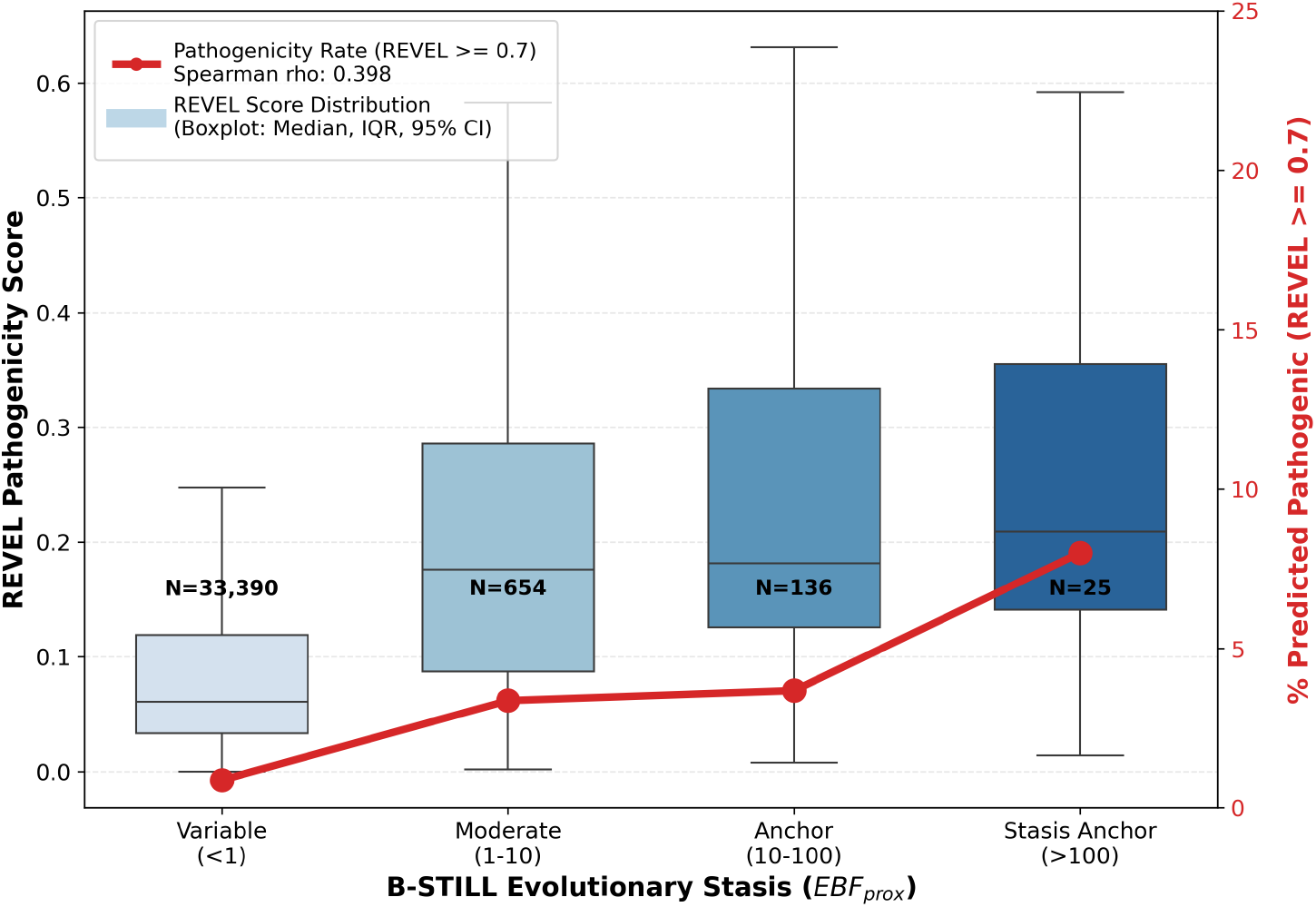
Genome-scale concordance between B-STILL inferred evolutionary stasis and REVEL pathogenicity scores (*N* = 34, 206 positions). The X-axis groups variants into four discrete stasis categories based on site-level *EBF*_*prox*_. The left Y-axis (blue boxplots) shows the distribution of continuous REVEL scores for each category, with the sample size (*N*) indicated at the bottom. The right Y-axis (red line) tracks the pathogenicity rate, defined as the percentage of variants within each category exceeding the high-confidence REVEL threshold (≥ 0.70). *ρ* represents the Spearman rank correlation coefficient, demonstrating an alignment between deep-time sequence constraint and modern clinical consensus.

### Identifying Regional Footprints of Overlapping Reading Frames in Viral Genomes

To evaluate whether the spans of regional Stasis Clusters correspond to genome regions that are expected to evolve under severe constraints, we applied the Hypergeometric Scan Statistic to the 110-gene FRESCO viral dataset (Sealfon <u>et al.</u>, 2015), where viral genomes were analyzed for the presence of overlapping reading frames. We utilized a significance threshold of *EBF* ≥ 10 to define the proximal ESAs, a choice supported by simulation results that demonstrate that this threshold maintains a Null FPR ≪ 1%. Application of the scan statistic identified 45 statistically significant regional Stasis Clusters (Table 4). B-STILL footprints documented overlapping reading frames across diverse viral families (Figure 7). In Hepatitis E Virus (ORF2), B-STILL identified a Stasis Cluster spanning 122 codons (sites 2–123, *p* = 2.7 × 10^−16^) that encompasses the 110-codon overlap with ORF3 (sites 15–124). Similar footprints were identified for the protein F frameshift in Hepatitis C (HCV1a sites 2–214, *p* = 4.4 × 10^−20^), the NSP6 overlap in Rotavirus A (NSP5 sites 2–98, *p* = 1.5 × 10^−7^), and the PIPO overlap in Turnip Mosaic Virus (polyprotein sites 983–1033, *p* = 1.7 × 10^−14^).

**Table 4:**
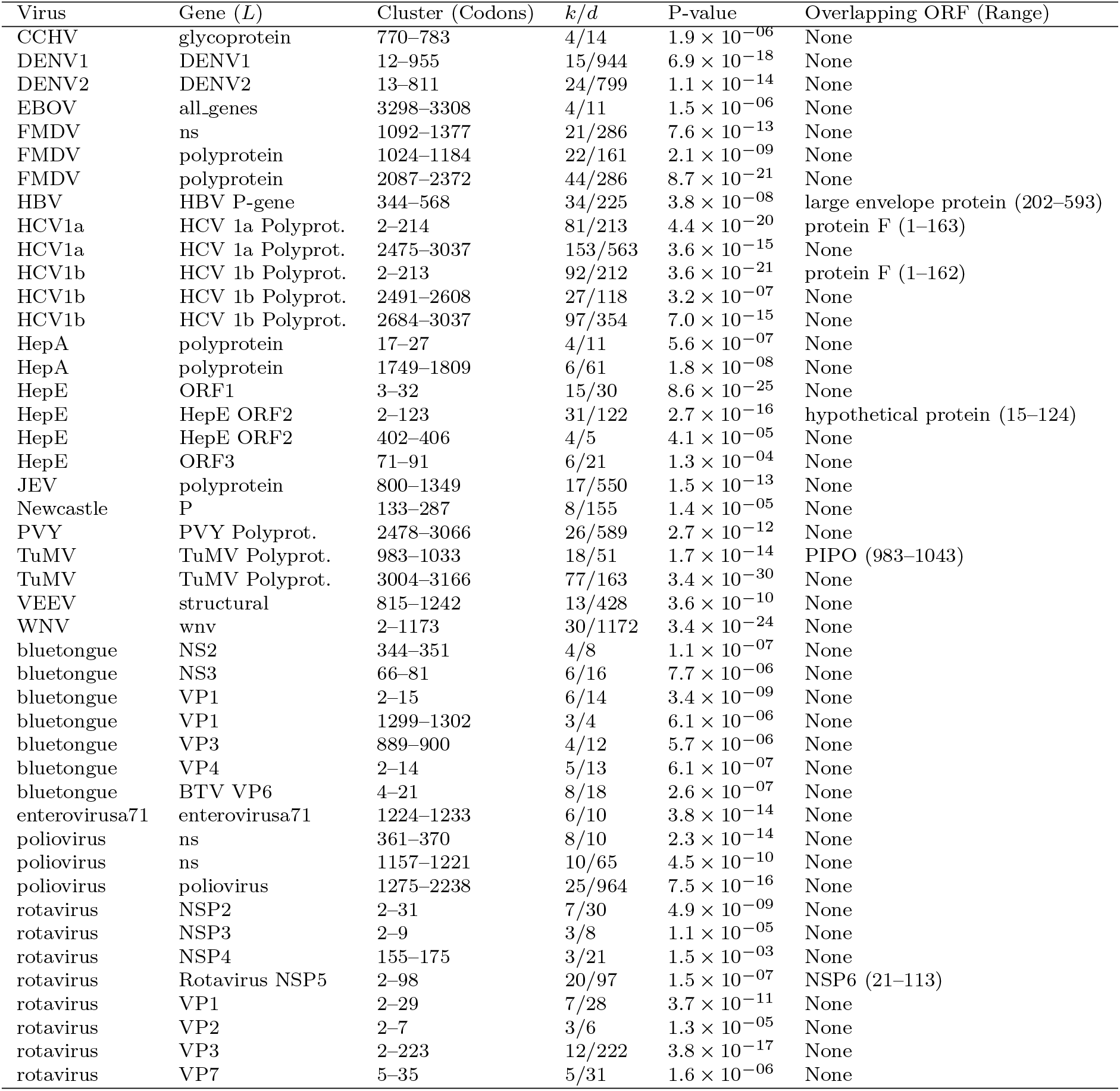
Statistically significant Stasis Clusters identified by Hypergeometric scan statistic (10,000 permutations) on FRESCO viral alignments. *k* represents stasis sites (*EBF* ≥ 10) in cluster span *d*.

**Figure 7:**
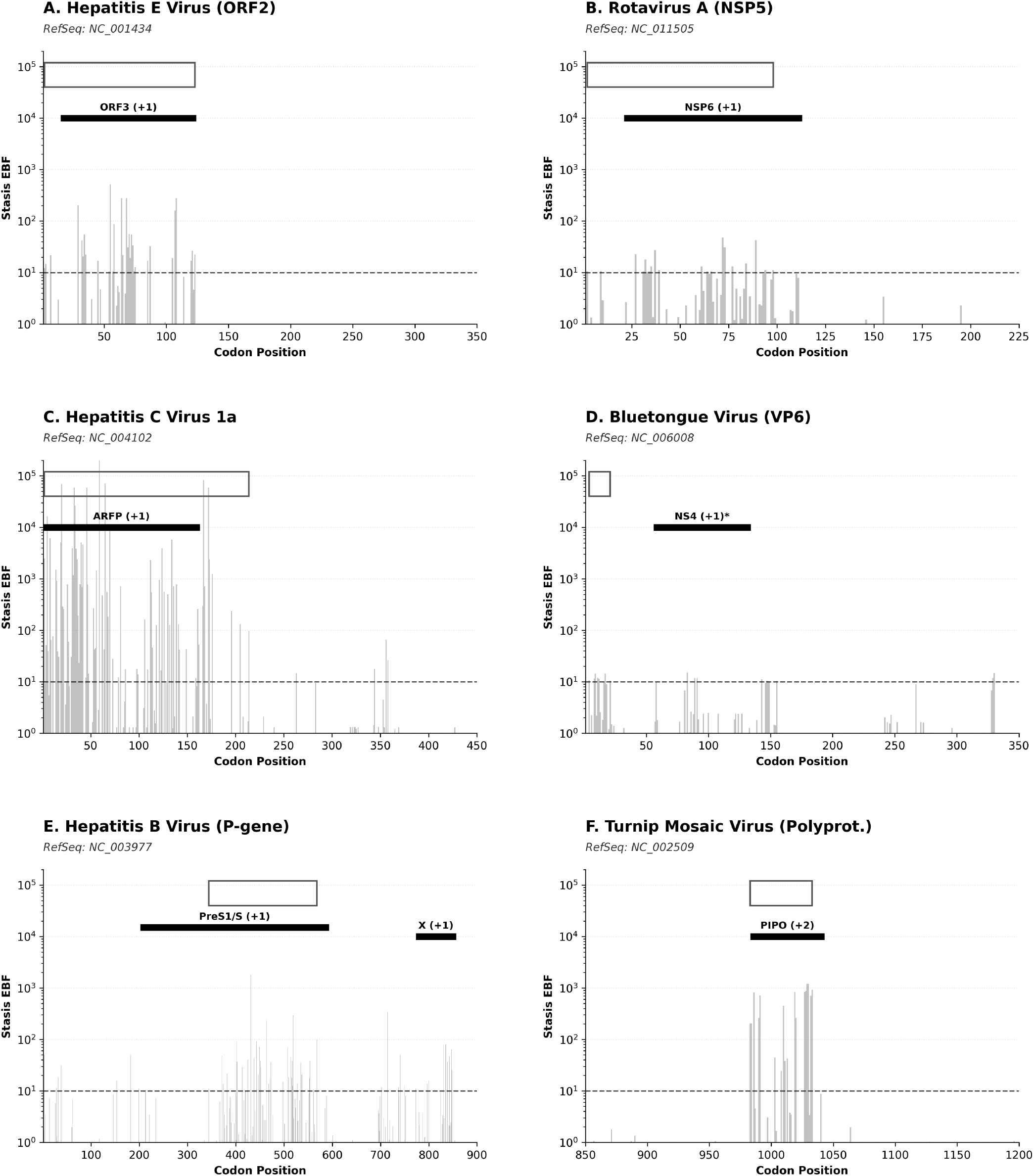
Mapping of B-STILL Stasis Clusters to annotated viral gene overlaps. Horizontal tracks indicate RefSeq overlaps (black lines) and statistically significant Stasis Clusters (dark gray rectangles).

However, detection of overlapping reading frames is not exhaustive. Of the 110 viral genes analyzed, several annotated overlaps remained undetected, particularly in cases where the primary or secondary reading frames exhibited low background mutational opportunity or relatively relaxed selection. This suggests that B-STILL targets the subset of features under significant purifying selection, in the context of a sufficiently variable genomic background. Furthermore, we identified numerous Stasis Clusters that do not correspond to annotated overlapping ORFs (Table 4). These “unmapped” regional Stasis Cluster signals likely footprint other multi-layered functional elements, such as structured cis-regulatory RNA motifs, viral packaging signals or biologically relevant nucleic acid secondary structural elements.

### Stasis Clusters in Mammals and the Dark Proteome

The global scan of the mammalian exome identified 4,888 statistically significant Stasis Clusters across 19,152 genes (Figure 9). The identified clusters exhibit a median span of 52 codons and are non-uniformly distributed across the exome, with ESA density peaking in genes involved in core cellular processes. Formal enrichment analysis using Fisher’s Exact Test against the InterPro database Blum <u>et al.</u> (2025) identified 133 functional associations (Table 5). We identified a significant enrichment of Stasis Clusters within genes associated with RNA-binding (OR = 2.4, *p <* 10^−15^), DNA-templated transcription (OR = 1.8, *p <* 10^−12^), and structural stabilization of the cytoskeleton (OR = 1.6, *p <* 10^−8^). Individual Stasis Clusters are non-randomly localized within critical protein architectures, including footprinting of the DNA-binding domain in `TP53`, the Armadillo repeat regions in `APC` and `NF1`, and numerous zinc finger motifs. These results demonstrate that B-STILL identifies regional selective footprints that correspond to the operational core of the proteome.

**Table 5:**
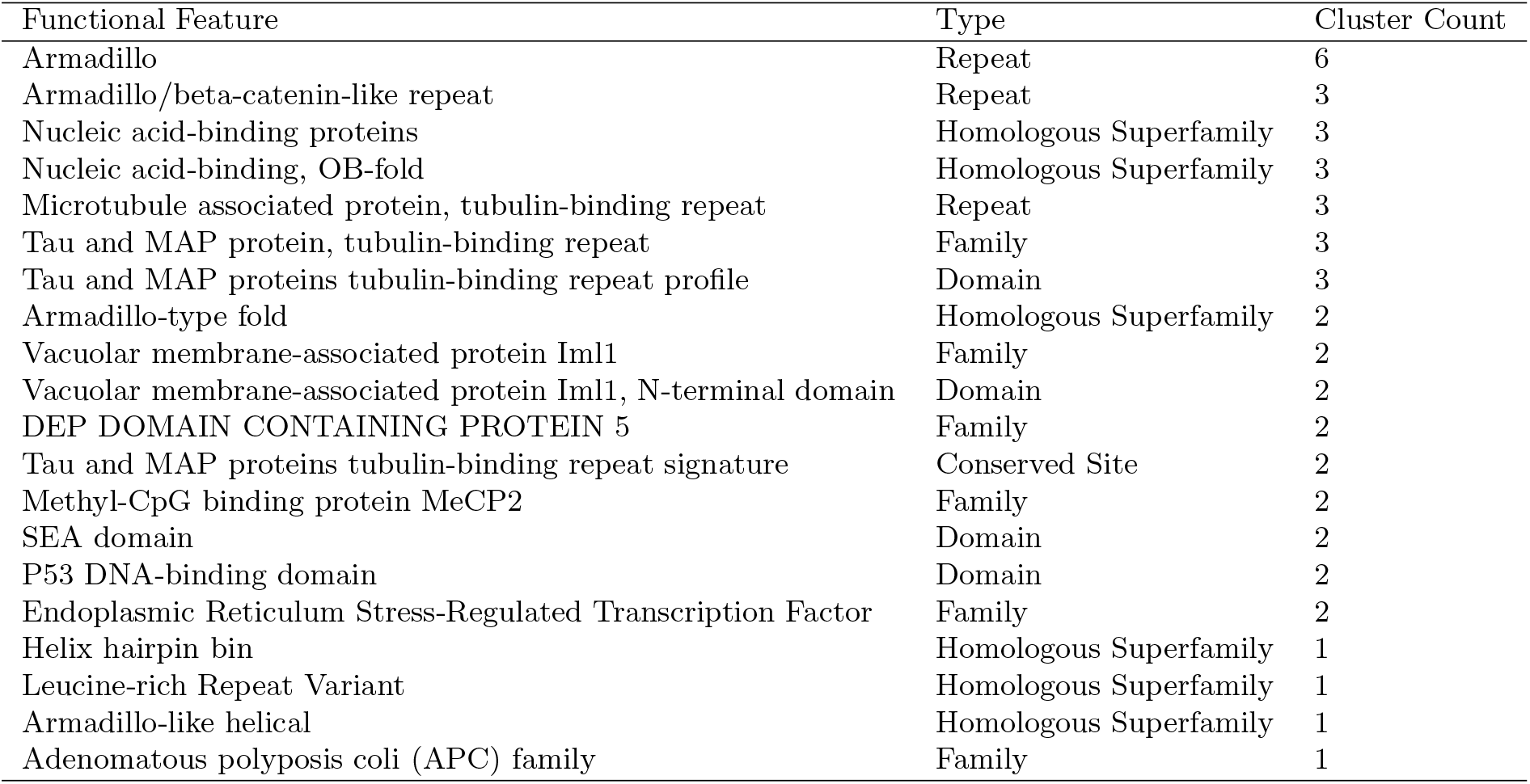
Top 20 functional protein features significantly enriched for B-STILL Stasis Clusters across the mammalian exome. Enrichment scores were calculated relative to the genomic background via Fisher’s Exact Test; all listed features exhibit *p <* 10^−50^ after multiple testing correction.

| Functional Feature | Type | Cluster Count |
| --- | --- | --- |
| Armadillo | Repeat | 6 |
| Armadillo/beta-catenin-like repeat | Repeat | 3 |
| Nucleic acid-binding proteins | Homologous Superfamily | 3 |
| Nucleic acid-binding, OB-fold | Homologous Superfamily | 3 |
| Microtubule associated protein, tubulin-binding repeat | Repeat | 3 |
| Tau and MAP protein, tubulin-binding repeat | Family | 3 |
| Tau and MAP proteins tubulin-binding repeat profile | Domain | 3 |
| Armadillo-type fold | Homologous Superfamily | 2 |
| Vacuolar membrane-associated protein Iml1 | Family | 2 |
| Vacuolar membrane-associated protein Iml1, N-terminal domain | Domain | 2 |
| DEP DOMAIN CONTAINING PROTEIN 5 | Family | 2 |
| Tau and MAP proteins tubulin-binding repeat signature | Conserved Site | 2 |
| Methyl-CpG binding protein MeCP2 | Family | 2 |
| SEA domain | Domain | 2 |
| P53 DNA-binding domain | Domain | 2 |
| Endoplasmic Reticulum Stress-Regulated Transcription Factor | Family | 2 |
| Helix hairpin bin | Homologous Superfamily | 1 |
| Leucine-rich Repeat Variant | Homologous Superfamily | 1 |
| Armadillo-like helical | Homologous Superfamily | 1 |
| Adenomatous polyposis coli (APC) family | Family | 1 |

**Figure 8:**
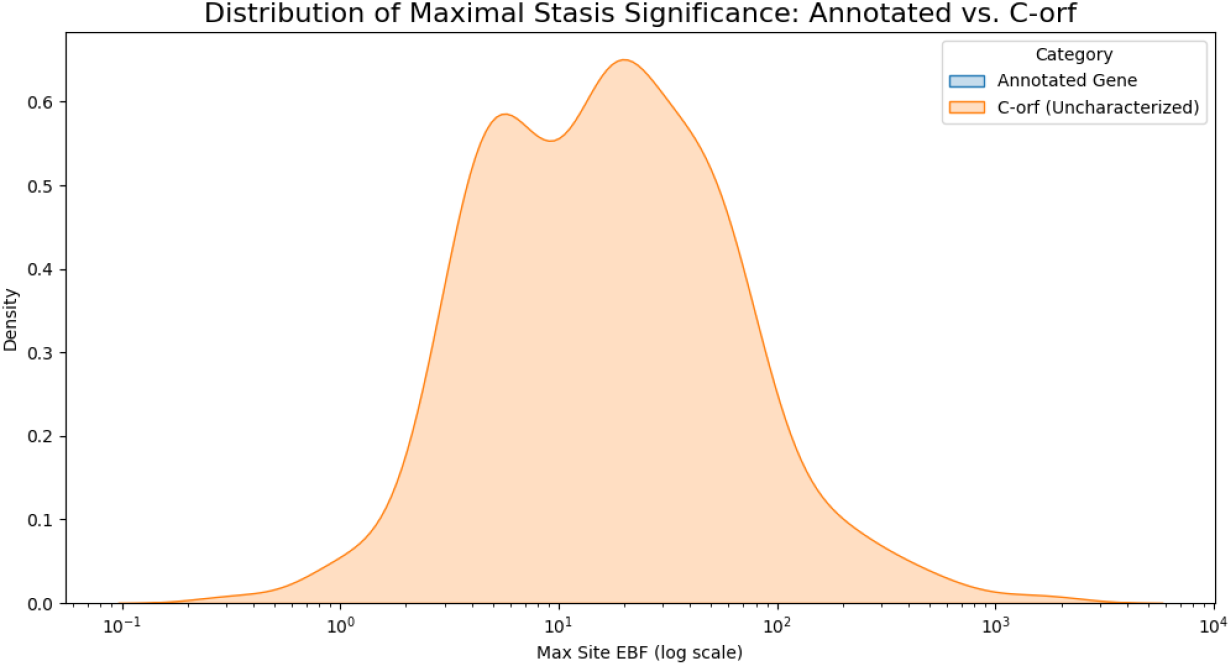
Distribution of Evolutionary Stasis Anchor significance (EBF) across 815 uncharacterized mammalian ORFs. The heavy-tailed distribution highlights a subset of sites under extreme constraint in the dark proteome.

**Figure 9:**
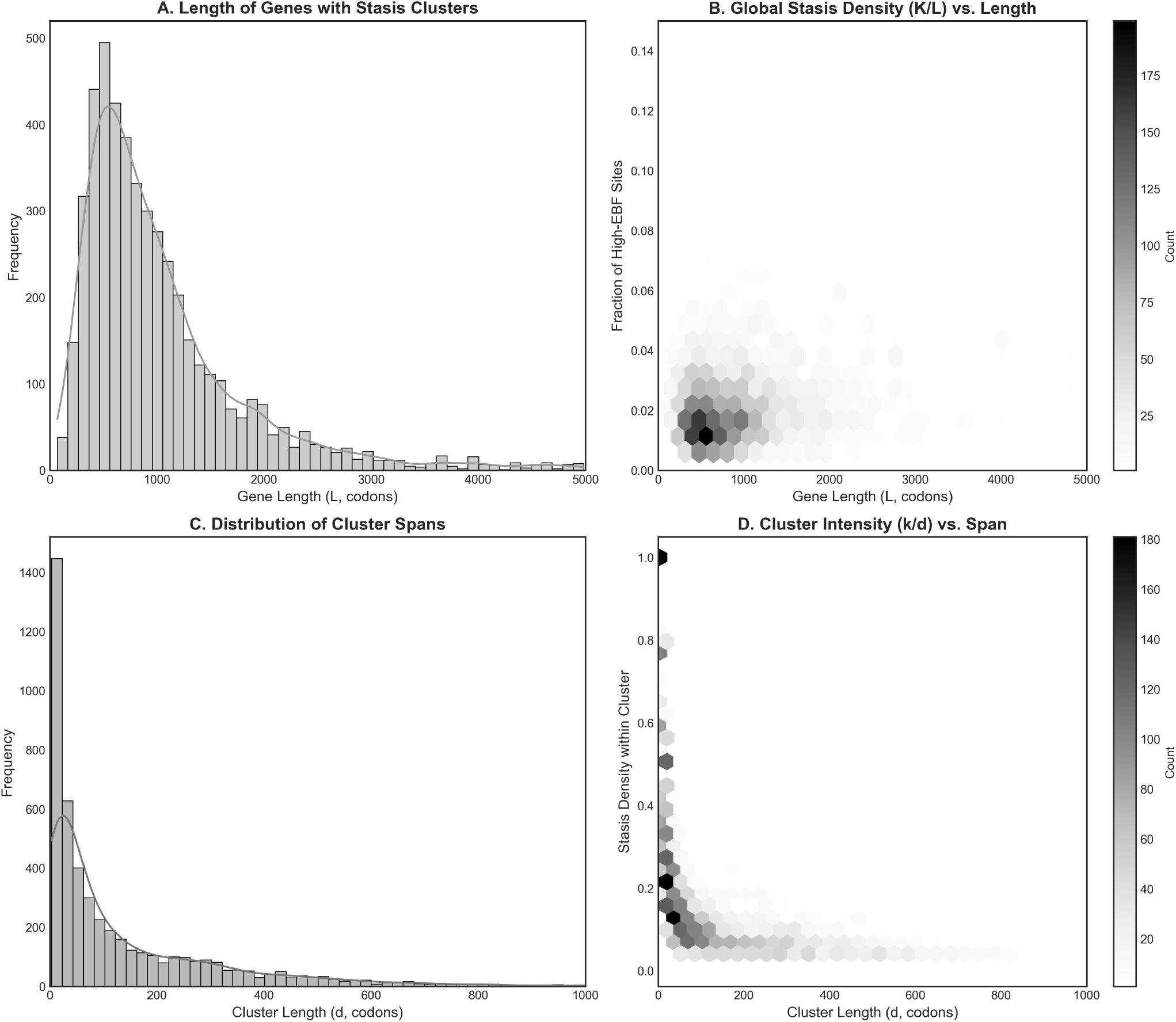
Statistical characterization of 4,888 Evolutionary Stasis Clusters identified across 19,152 mammalian genes.

Application of the B-STILL framework to 815 uncharacterized protein-coding genes (the “dark proteome”) of the 120-species mammalian alignment identified 1,342 ESAs and eight statistically significant regional Stasis Clusters (Figure 8). In these proteins of unknown function, the discovery of contiguous ESAs provides a data-driven approach to identifying functional modules.

A particularly notable case is found in `FAM214A`, which contains a 239-codon Stasis Cluster (*p* = 1.4 × 10^−18^, sites 822–1060) harboring multiple ESAs (*EBF >* 500). Statistical analysis of the `FAM214A` structural model reveals that these signals are statistically significantly clustered in three-dimensional space (*p <* 10^−4^, permutation test; Figure 10). We identified a structural hub centered at residue 1038, which contains nine ESAs within a 10 Å radius. Similarly, we identified a 1,812-codon cluster in `C2ORF16` (*p* = 8.3 × 10^−14^) and a small 3-residue Stasis Cluster in `C1ORF167` (*p* = 6.9 × 10^−7^, sites 209–211). These results demonstrate that regional Stasis Clusters can pinpoint the structural frameworks or interaction interfaces of uncharacterized proteins, providing targets for experimental functional characterization.

**Figure 10:**
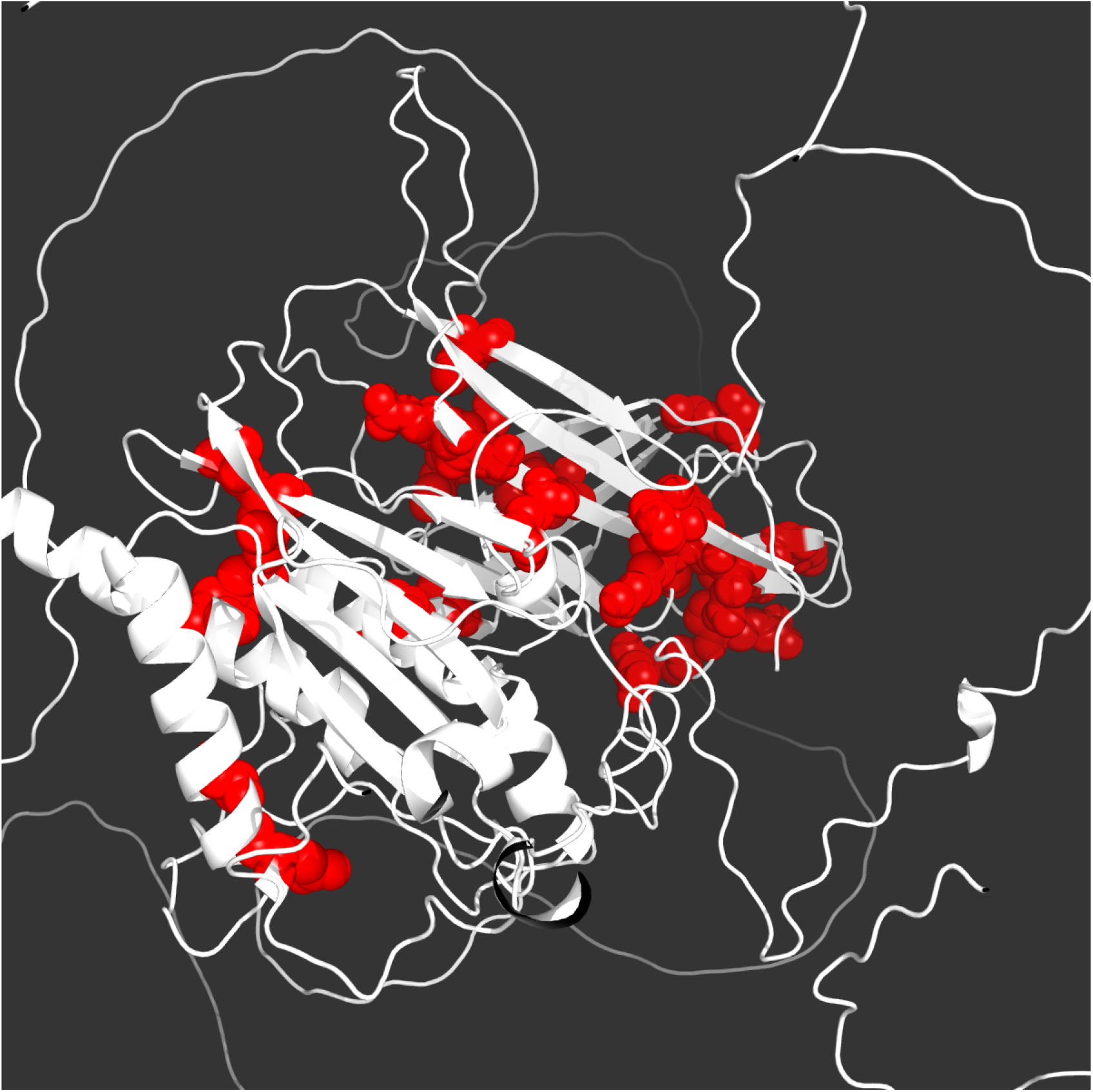
Three-dimensional structural hub of Evolutionary Stasis Anchors in the uncharacterized protein FAM214A. Residues are colored by B-STILL significance (log_10_ *EBF*), with significant anchors (*EBF* ≥ 100) shown as red spheres. We identified a significant Stasis Cluster comprising nine Evolutionary Stasis Anchors centered at residue 1038 (*p <* 10^−4^, permutation test), likely demarcating a structural or interaction hub in this poorly characterized ORF.

### Extremes of the Mammalian Evolutionary Stasis Landscape

We identified a subset of genes harboring long contiguous Stasis Clusters spanning more than 1,000 codons. The most prominent example is found at the C-terminus of `MUC16`, where a 2,063-codon cluster (*p* = 1.8 × 10^−24^) precisely footprints a series of 16 tandemly repeated SEA domains. While alignment inaccuracies typically increase apparent variation, B-STILL identifies contiguous blocks of zero-variation, suggesting that the stasis signal is a biological property of these regions. Similar long Stasis Clusters were identified in `NEB` (Nebulin, 1,193 codons), where absolute sequence constraint maintains the precise molecular spacing required for muscle thin-filament regulation. Conversely, the high-intensity tail of the distribution identifies “functional snaps”: short, contiguous intervals of absolute or near-absolute sequence stasis (*k/d* ≥ 0.80) where nearly every codon is an ESA. These motifs frequently map to critical catalytic or binding interfaces (Table 5). For instance, we identified a 6-residue snap in `KPNB1` (Importin *β*-1, sites 38–43, *p* = 1.6 × 10^−8^) mapping to a structural ARM repeat, and a 7-residue snap in `SIM2` mapping to a functional PAC motif (*p* = 4.8 × 10^−12^). Other notable snaps include a 6-residue Stasis Cluster in the Rab GTPase-activating domain of `EVI5L` (*p* = 7.5 × 10^−14^) and a 3-residue cluster in the RNA polymerase II clamp domain (`POLR2A`, sites 11–13).

### Comparative Resolution of B-STILL vs. phyloP

Head-to-head benchmarking against phyloP across 38 representative mammalian genes demonstrates that B-STILL provides higher resolution for functional ESAs than standard nucleotide-based LRTs (Figure 11). While there is a general correlation between metrics, the relationship is stronger when comparing B-STILL Proximal EBFs to the minimum (worst) phyloP score within a codon (*ρ* = 0.35) rather than the maximum (*ρ* = 0.19). This result highlights a distinction in how these models operate: because standard nucleotide-based methods treat sites independently, they often assign high significance to first or second codon positions that are invariant due to intense purifying selection on the encoded amino acid. This creates a “0/0 plateau” where frequentist scores saturate at a significance ceiling; because the LRT is a function of the observed data, all invariant sites across a phylogeny receive an identical score regardless of their underlying mutational opportunity. B-STILL, by modeling the codon as the fundamental unit of selection and leveraging a hierarchical prior, breaks this resolution ceiling by ranking stasis according to its statistical surprise relative to the gene-wide background.

**Figure 11:**
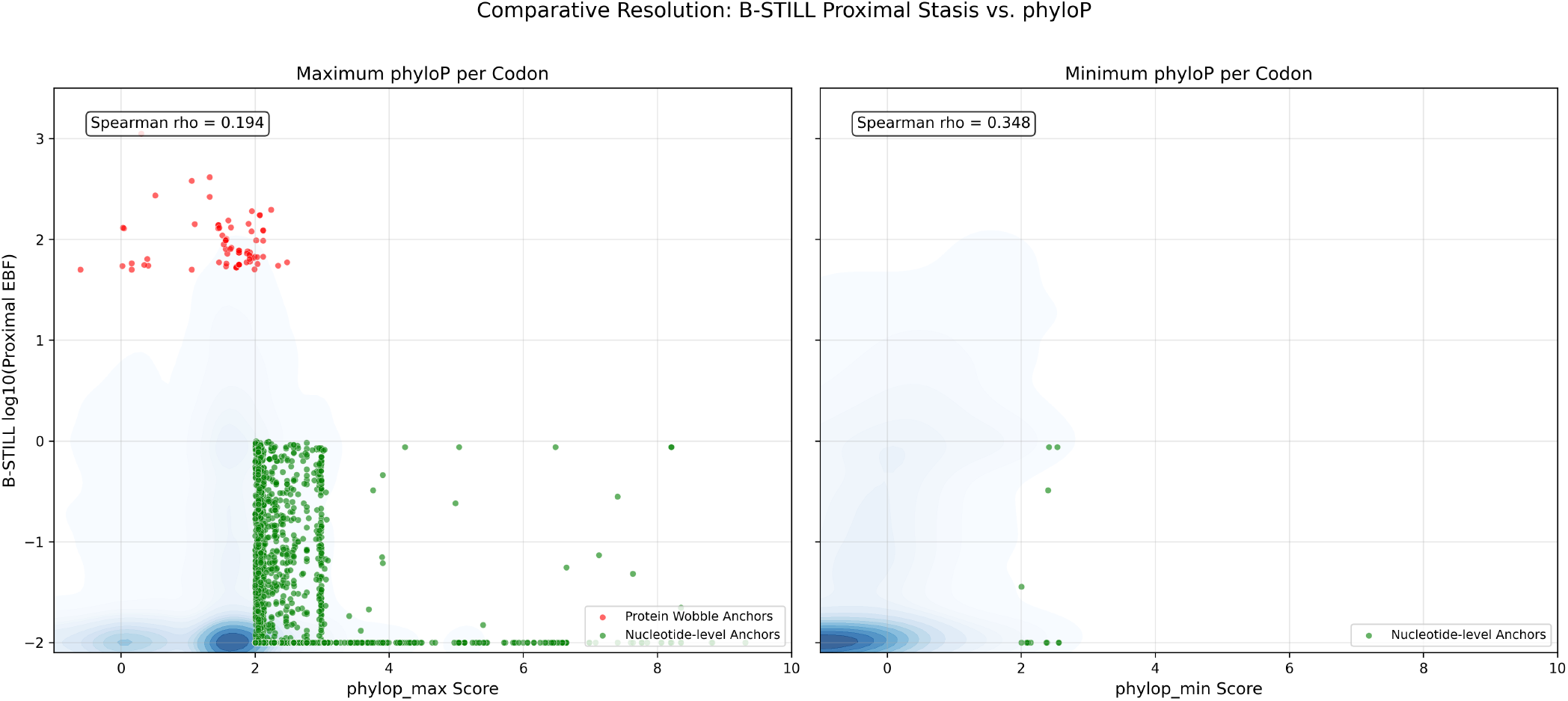
Comparative resolution of B-STILL Proximal Stasis EBF vs. nucleotide-level conservation (phyloP) across 38 mammalian genes. (Left) Correlation with the maximum phyloP score per codon (*ρ* = 0.19). The visible “0/0 plateau” represents the resolution ceiling of frequentist Likelihood Ratio Tests (LRTs), which assign an identical significance score to all invariant sites, failing to distinguish between stochastic invariance and functional constraint. (Right) Correlation with the minimum phyloP score per codon (*ρ* = 0.35). The stronger alignment with the least-conserved position indicates that B-STILL EBFs effectively filter out genetic-code artifacts—where positions are invariant solely due to codon-structure constraints—identifying functional anchors with higher specificity.

This distinction is illustrated by two emergent archetypes of conservation. The first, termed “Protein Wobble Anchors,” consists of sites where B-STILL identifies stasis (Proximal EBF *>* 100) while the maximum phyloP score remains low (*<* 3.0). At site 59 of `APOBEC3D`, for instance, B-STILL reports a Proximal EBF of 1124.3, while the maximum phyloP score is 0.30. Similarly, at site 2103 of `F8` (Proximal EBF = 272.8, max phyloP = 0.51) and site 2777 of `BRCA2` (Proximal EBF = 381.0, max phyloP = 1.05), B-STILL identifies purifying selection that is not captured by nucleotide-level tools. In these cases, the protein sequence is invariant across 120 mammals, but the underlying codons exhibit synonymous variation (e.g., `GTA` vs. `GTG` at `APOBEC3D` site 59). Standard nucleotide-based methods conflate this synonymous variation with a lack of selective constraint, resulting in lower scores. B-STILL identifies that the absence of amino-acid substitutions in the face of background synonymous opportunity is a signature of functional constraint.

Conversely, the comparison identifies “Nucleotide-Level Anchors,” where phyloP scores are high but B-STILL Proximal EBFs remain low. These sites often correspond to positions where both synonymous and non-synonymous substitutions are rare at the nucleotide level, but B-STILL identifies amino-acid variation that suggests the position is not evolutionarily invariant at the protein level. For instance, at site 5 of `TP53`, phyloP yields a score of 2.68, whereas B-STILL assigns a negligible Proximal EBF (7.6 × 10^−8^). Alignment analysis reveals that while the consensus `CAG` (Glutamine) is prevalent, multiple lineages exhibit non-synonymous transitions to Proline (`CCG`) and Leucine (`CTG`), identifying that the site is not a strict functional anchor.

### Computational Performance

Benchmarks across 19,152 genes demonstrate B-STILL’s high-throughput performance (Table 6), completing the analysis of nearly 30 million codons in approximately 422 CPU-hours on a cluster of Ampere Altra (3.0 GHz) ARM64 compute nodes. The framework achieved a mean throughput of 19.52 codons per second/per core, establishing it as a scalable tool for high-resolution genomic footprinting.

**Table 6:** Computational performance of the B-STILL framework across the 120-species mammalian exome dataset. Benchmarks were recorded on a cluster of Ampere Altra (3.0 GHz) ARM64 compute nodes.

| Metric | Value |
| --- | --- |
| Total genes analyzed | 19,152 |
| Total codons analyzed | 29,691,139 |
| Mean overall runtime per gene | 79.4 s |
| Median overall runtime per gene | 74.0 s |
| Mean throughput | 19.52 codons/s |
| Total CPU-hours (Ampere Altra 3.0 GHz) | $\approx 422$ h |

## Discussion

B-STILL provides a principled Bayesian framework for quantifying selective and evolutionary constraint. A central conceptual advancement of this work is the surmounting of the “0/0 plateau” that limits the standard frequentist tools that are commonly used to quantify conservation and purifying selection. Because Likelihood Ratio Tests are functions of the observed data, they exhibit a resolution ceiling where all invariant sites in a given phylogeny receive an identical maximal score. By conditioning the stasis signal on site-specific synonymous evolutionary opportunity and gene-wide selective priors, B-STILL breaks this degeneracy, providing a ranking of evolutionary stasis based on statistical surprise. An example of this are “Protein Wobble Anchors”—sites where extreme evolutionary stasis persists despite high synonymous substitution opportunity. This hierarchical design effectively filters out the global selective background: while an invariant site in an ultra-conserved gene (e.g., a histone) is expected and thus yields a moderate EBF, a single invariant anchor in a rapidly diversifying gene (e.g., an olfactory receptor) represents a profound deviation from the neutral expectation and is prioritized accordingly. This ensures that B-STILL identifies the “operational core” of a protein relative to its specific evolutionary context, rather than merely re-stating global conservation.

The framework’s codon-aware design provides an advantage for identifying regulatory stasis that persists at the nucleotide level independently of amino-acid change. Our validation against the ClinVar database demonstrates that Synonymous Stasis (*EBF*_*syn*_) is a predictor of protein sites at which any amino acid substitutions are likely to be clinically pathological (aggregate AUROC = 0.88); identifying critical nodes in genes like `TP53` and `CFTR` that are invisible to ensemble protein-focused predictors of clinical relevance such as REVEL. Interestingly, B-STILL’s predictive power for synonymous variants (AUROC = 0.88) exceeds its performance for missense variants (AUROC = 0.65). This “synonymous paradox” likely reflects the fact that the pathogenicity of synonymous substitutions is driven by relatively pure nucleotide-level constraints (e.g., splicing enhancers, mRNA stability) that are well-captured by a phylogenetic baseline. In contrast, missense pathogenicity is a composite of phylogenetic, structural, and biochemical features; by providing a highly specific phylogenetic prior, B-STILL complements rather than replaces modern ensemble methods, which often struggle to calibrate the “evolutionary floor” in the absence of protein-sequence change.

The integration of site-level ESAs into regional Stasis Clusters revealed larger-scale footprints of extreme evolutionary constraint across diverse genomic architectures. In viral genomes, B-STILL provides high-precision footprinting of overlapping reading frames, identifying the intervals where multi-layered coding requirements enforce absolute sequence immobilization. In the mammalian exome, these regional Stasis Clusters map to critical architectural features, including the SEA domain repeats in `MUC16`, the precise molecular spacing interfaces in `NEB`, and structural motifs such as the ARM repeats in `KPNB1`. The discovery of these clusters independently of structural or biochemical data suggests that regional stasis is a robust proxy to identify the absolutely indispensable functional modules of the proteome.

B-STILL is particularly suited for functional annotation within the “dark proteome.” In uncharacterized ORFs where biochemical data are absent, the spatial clustering of ESAs provides a data-driven strategy for identifying critical residues. The discovery of a statistically significant structural hub in `FAM214A` (*p <* 10^−4^) provides a prioritized target for experimental characterization, demonstrating that distribution across a gene of ESAs can demarcate potentially functional protein interfaces in the absence of any known homology or mechanistic biochemical data.

Finally, while the genome annotation field is increasingly moving toward phylogeny-aware genomic language models (gLMs) such as GPN-Star and AlphaGenome, B-STILL remains a tool for high-resolution, mechanistic-agnostic annotation. Unlike deep learning models that synthesize vast contextual windows through complex neural architectures, B-STILL provides a transparent, data-driven codon-level map of selective constraint derived directly from the underlying substitution process. By focusing on the genomic syntax of coding regions across intermediate to deep evolutionary timescales, B-STILL provides principled codon-level estimates of functionally relevant evolutionary stasis that complement, and could therefore also productively augment, modern AI-driven genome annotation tools.

## Acknowledgements

This work was supported in part by the National Institutes of Health (R01 AI183870, R01 GM151683) and the National Science Foundation (DBI 2419522).

## Data and Code Availability

B-STILL is implemented in HyPhy (v2.5.95+). Source code, analysis scripts, and simulation datasets are available under the MIT license at https://github.com/veg/hyphy and https://github.com/veg/b-still.

