## Supplementary Materials for "Beyond Invariable Sites: Using Evolutionary Stasis to Map Multi-Layered Constraints on the Evolution of Viral and Mammalian Genomes"

### Supplementary Material: A Technical Review of Methodologies for Estimating the Probability of Site-Level Conservation

Sergei L. Kosakovsky Pond Darren P. Martin

April 9, 2026

#### 1 Introduction

The quantification of site-specific conservation is a cornerstone of comparative genomics, yet the statistical machinery underlying various tools differs significantly in their assumptions regarding neutral drift and rate heterogeneity. To provide a rigorous context for the development of B-STILL, we dissect the technical mechanisms by which existing methods estimate the probability of conservation. Ignore the nuances of these implementations at your own peril, as each introduces distinct statistical trade-offs between sensitivity and specificity.

#### 2 Methodological Dissections

##### 2.1 phyloP: Phylogenetic P-values

The phyloP tool, part of the PHAST suite, represents perhaps the most direct application of likelihood ratio tests (LRT) to site-level conservation [2].

- Estimation Logic: For each site, phyloP evaluates the observed number of substitutions against a pre-calibrated neutral phylogenetic tree. The null hypothesis assumes that substitutions follow a neutral Markov process (typically calibrated using four-fold degenerate sites).
- Quantification of Invariance: For a strictly invariant site, the surprisingness is a direct function of the total neutral tree length ($T$). Under a Poisson approximation, the probability of zero substitutions is $e^{-T}$. phyloP outputs a score defined as $-\log_{10}(pext{-}value)$, where $p$ is the probability of the data under the neutral model.
- Limitation: As a site-independent method, it lacks the ability to share information across the gene, making it highly sensitive to the accuracy of the provided neutral tree.

##### 2.2 GERP++: Genomic Evolutionary Rate Profiling

GERP++ eschews p-values in favor of a count-based metric known as Rejected Substitutions (RS) [1].

- Estimation Logic: The method calculates the expected number of substitutions ($E$) that would have occurred at a site under neutral evolution given the tree topology and branch lengths. It then subtracts the observed number of substitutions ($O$).

- Quantification of Invariance: For an invariant site ($O = 0$), the RS score is exactly equal to the neutral tree length ($RS = E$). This provides an intuitive, additive measure of constraint: an RS score of 5.0 implies that evolution has successfully "rejected" five expected mutations.
- Limitation: Like phyloP, GERP++ treats sites in isolation, failing to account for the gene-wide distribution of selective pressures.

##### 2.3 phastCons: Spatial Context via HMMs

Unlike independent site methods, phastCons utilizes a Phylogenetic Hidden Markov Model (phylo-HMM) to contextualize conservation [**?**].

- Estimation Logic: The model toggles between two hidden states: "neutral" and "conserved." The state transitions are governed by a set of parameters that reflect the expected length of conserved elements.
- Quantification of Invariance: It assigns a posterior probability (from 0 to 1) that a site belongs to the conserved state. A strictly invariant site flanked by high variation will receive a lower probability than one residing within a cluster of conserved positions.
- Limitation: While effective for discovering conserved motifs, the smoothing effect of the HMM can obscure sharp, site-specific signatures of purifying selection.

##### 2.4 Rate4Site: Empirical Bayesian Rate Inference

For amino acid alignments, Rate4Site is the established gold standard for quantifying protein-level conservation [**?**].

- Estimation Logic: The method uses Empirical Bayesian inference to assign a relative evolutionary rate to every site, incorporating a substitution matrix (e.g., WAG or JTT) and the phylogeny.
- Quantification of Invariance: An invariant site yields a posterior probability distribution sharply peaked at the lowest possible rate. This allows for a continuous ranking of sites even when multiple sites appear identical at the sequence level.
- Limitation: It typically assumes a single global rate for the entire protein, which is fundamentally insufficient for capturing the $dN/dS$ dynamics essential for codon-level analysis.

##### 2.5 Mechanistic Codon Models (HyPhy / PAML)

Codon-based models provide a more nuanced view of selection by estimating the ratio of non-synonymous to synonymous substitutions ($\omega = \beta/\alpha$) [**?**, **?**].

- Estimation Logic: These models use synonymous drift ($\alpha$) as an internal control. Conservation is inferred when $\beta < \alpha$.
- Quantification of Invariance: Methods like FEL or FUBAR estimate the posterior probability $P(\beta < \alpha | extData)$. However, at strictly invariant sites where $\alpha = \beta = 0$, the standard selection test is often statistically uninformative.
- Conceptual Gap: This methodological failure at the limit of zero substitutions is the primary motivation for the high-resolution grid and EBF approach implemented in B-STILL.

##### 2.6 EVE: Generative Deep Learning

The EVE (Evolutionary model of Variant Effect) method utilizes Variational Autoencoders (VAEs) trained on deep sequence alignments [?].

- Estimation Logic: EVE learns the latent multi-dimensional distribution of sequences. It quantifies conservation through the "unlikeliness" of a mutation within the learned manifold.
- Quantification of Invariance: For invariant sites, the model's latent space dictates that any amino acid other than the consensus has an extremely low probability.
- Limitation: While powerful, these models require large-scale alignments and are often computationally prohibitive compared to grid-based Bayesian methods.

#### 3 Standardizing the Neutral Baseline for Benchmarking

To rigorously compare B-STILL against site-independent tools like phyloP and GERP++, it is essential to establish a conceptually equivalent neutral baseline. Traditionally, these tools rely on neutral trees calibrated using four-fold degenerate sites—an approximation that is fundamentally insufficient as it discards the majority of the synonymous signal and introduces potential compositional biases.

We propose a Codon-derived Neutrality protocol for benchmarking:

1. Synonymous Tree Estimation: Utilize a mechanistic codon model (e.g., MG94) to estimate the synonymous substitution rate ($\alpha$) across the entire alignment. This leverages every synonymous substitution opportunity, yielding a significantly more stable and data-dense phylogeny than 4-fold site counting.
2. Neutral Reference Extraction: The true neutral baseline for a coding region is defined as the tree where branch lengths ($L$) represent expected synonymous change ($L_{neutral} = \alpha \times T$). This tree represents the expected drift at a site where $\beta = \alpha$.
3. Cross-Tool Calibration: Export this synonymous tree and the associated GTR substitution parameters into a PHAST-compatible `.mod` file. This ensures that the "surprise" measured by phyloP is statistically comparable to the constraint inferred by B-STILL, moving the comparison from one of data-proxies to one of statistical inference frameworks.
